# A CRISPR toolbox for generating intersectional genetic mice for functional, molecular, and anatomical circuit mapping

**DOI:** 10.1101/2021.06.10.447908

**Authors:** Savannah J. Lusk, Andrew McKinney, Patrick J. Hunt, Paul G. Fahey, Jay Patel, Andersen Chang, Jenny J. Sun, Vena K. Martinez, Ping Jun Zhu, Jeremy R. Egbert, Genevera Allen, Xiaolong Jiang, Benjamin R. Arenkiel, Andreas S. Tolias, Mauro Costa-Mattioli, Russell S. Ray

## Abstract

**Background:** A full understanding of circuits and cellular mechanisms governing health and disease requires the dissection and multi-faceted study of discrete cell subtypes in developing and adult animal models. Recombinase-driven expression of transgenic response alleles represents a significant and powerful approach to delineate cell populations for functional, molecular, and anatomical study. In addition to single recombinase systems, the expression of two recombinases in distinct, but partially overlapping, populations allow for more defined target expression. Although the application of this method is becoming increasingly popular, the expense and difficulty associated with production of customized intersectional mouse lines have limited widespread application to more common allele manipulations that are often commercially produced at great expense.

**Results:** We present a simplified CRISPR toolkit for rapid, inexpensive, and facile intersectional allele production. Briefly, we produced 7 intersectional mouse lines using a dual recombinase system, one mouse line with a single recombinase system, and three embryonic stem (ES) cell lines that are designed to study how functional, molecular, and anatomical features relate to each other in building circuits that underlie physiology and behavior. As a proof-of-principle, we applied three of these lines to different neuronal populations for anatomical mapping and functional *in vivo* investigation of respiratory control. We also generated a mouse line with a single recombinase-responsive allele that controls the expression of the calcium sensor Twitch-2B. This mouse line was applied globally to study the effects of follicle stimulating hormone (FSH) and luteinizing hormone (LH) on calcium release in the ovarian follicle.

**Conclusions:** Lines presented here are representative examples of outcomes possible with the successful application of our genetic toolkit for the facile development of diverse, modifiable animal models. This toolkit will allow labs to create single or dual recombinase effector lines easily for any cell population or subpopulation of interest when paired with the appropriate Cre and FLP recombinase mouse lines or viral vectors. We have made our tools and derivative intersectional mouse and ES cell lines openly available for non-commercial use through publicly curated repositories for plasmid DNA, ES cells, and transgenic mouse lines.

## BACKGROUND

Targeted expression of effector molecules, like fluorescent markers, calcium reporters, optogenetic actuators, or exogenous ligand-responsive receptors (DREADDs) (1), are increasingly applied in a variety of fields for greater precision, quantitative expression level control, and reduced side effects compared to previous methods for labeling and manipulation. For example, the Cre/LoxP system is used to achieve permanent cell-type restriction by using a promoter or enhancer to express the site-specific recombinase Cre (2). In such examples, Cre transgenes are paired with a constitutively active, but conditional, allele where expression of an effector molecule, such as eGFP, is interrupted by a LoxP-Stop-LoxP cassette that is recombined out by Cre, which enables expression in targeted, Cre-expressing cells. The use of various effector molecules in this paradigm enables fluorescent marking, neuronal perturbation, molecular affinity pull-downs, activity tracking, and other studies (1,3–28).

In many fields, it is becoming increasingly clear that recombinase expression based on a single gene does not offer the resolution needed for a variety of developmental or targeting applications (29–31). Indeed, application of intersectional genetics has led to new progress in various fields including neural circuits, cell type lineage, and embryonic development (32–34). Intersectional genetics adds needed resolution by employing a dual recombinase system using both Cre recombinase as well as a second recombinase, FLP, to activate a conditional effector allele only in cells where both recombinases have been expressed in the same cell (though not necessarily concurrently) (20,35–42). With new methodologies being developed to use Cre and FLP not only as traditional genetic markers, but also as activity and connectivity markers, unique combinatorial cell type definitions become possible (43–47). Thus, a resource consisting of multiple dual recombinase intersectional alleles that each express different effector molecules would add significant value and needed resolution to our ability to deconstruct neural circuits on multiple levels. This technology could also be applied to a multitude of other fields where intersectionally defined subpopulations of target cells exist and may play different roles in the measured outcomes.

Although mouse intersectional technology provides relatively benign access to otherwise inaccessible populations of cells, few laboratories have generated single transgenic or intersectional genetic mouse lines in house for several reasons. The complexity and size of the final targeting vectors puts them beyond present (cost-effective) commercial DNA synthesis capabilities, thus requiring some level of recombinant DNA cloning and precluding straightforward production of large, high fidelity ssDNA donors that can facilitate pro-nuclear CRISPR-mediated targeting (48). While intersectional targeting vectors are available from the Addgene plasmid repository, they are finished vectors that require significant reverse engineering or modifications for use in a new context or approach. To our knowledge, there are no modular intersectional targeting vectors that are publicly available for facile and rapid production of new targeting alleles for the generation of intersectional mouse models. Furthermore, vector stability and other *in vitro* difficulties combined with the expense and time associated with target vector insertion and mouse line production limit the number of intersectional mouse lines available for public use. Thus, the production of intersectional genetic mouse lines has been largely limited to a small number of pioneering labs or resource rich institutions such as the Howard Hughes Medical Institute and the Allen Institute for Brain Science (41, 49).

To address these pitfalls and make the production of intersectional genetic mouse models more widely feasible, we aimed to produce a freely available resource toolbox consisting of several intersectional and single-recombinase responsive *Rosa26* targeting vectors for rapid, facile, and cost-effective generation of complex mouse lines using CRISPR/Cas9-mediated homologous recombination in mouse embryonic stem cells and early oocytes. In oocytes, genomic insertions/deletions (in/dels) and short targeted insertions are readily produced whereas large construct targeting in oocytes has shown more limited but growing success. Additionally, requisite equipment and facilities are difficult to access and out of reach for many investigators. ES cell approaches are well established and widely available, allowing for rapid screening and identification, and, if using morula aggregation, require much simpler methodologies and equipment for mouse generation.

Towards this, we produced 7 intersectional mouse lines using a dual recombinase system; one mouse line with a single recombinase system, and three additional ES cell lines to study how functional, molecular, and anatomical features relate to each other in building the circuits that underlie physiology and behavior. As a proof of principle, we applied three of these lines to different neuronal populations for anatomical mapping and functional *in vivo* characterization in respiratory control. Next, we globally applied the single recombinase-responsive line, which controls the expression of the calcium sensor Twitch-2B. Twitch-2B was expressed globally in the generated mouse line to study the effects of follicle stimulating hormone (FSH) and luteinizing hormone (LH) on calcium release in the ovarian follicle. The publication and availability of this technology will allow for the seamless production of a highly diverse group of mouse lines that can be used to generate animal models of human disease, label specific cell populations for developmental or connectivity studies, or modulate cellular activity in established disease model lines, among other possibilities. All reagents and vectors used or generated in this study are now openly available for not-for-profit research.

## RESULTS

### Vector design and optimization

For each of the targeting vectors generated, the intersectional or Dre-responsive cassette was knocked into a well-established site in the *Rosa26* locus (**Fig. 1)** (50, 51). For positive ES cell clone selection, the targeting vector was simplified and improved in several ways compared to earlier approaches (52). A simplified neomycin resistance cassette was integrated into the intersectional cassette before the second LoxP-flanked stop cassette to utilize the *CAG* promoter and polyadenylation (*pA*) sequences of the FRT flanked stop cassette, eliminating prior use of an additional PGK promoter and *Bovine Growth Hormone polyA* (BGHpA) signal. To use CRISPR/Cas9, we cloned an sgRNA that was close to the 5’/3’ homology junction into the px330 vector (53), which expresses both Cas9 and the subcloned sgRNA (*px330_Rosa26_sgRNA*). A Woodchuck Hepatitis Virus (WHV) Posttranscriptional Regulatory Element (WPRE) and BGHpA were added at the end of the expression cassette to enhance effector molecule expression and limit reliance on the disrupted Rosa Locus for transcript termination (52). The homology arms were significantly shortened to 1Kb to remove repetitive genomic sequences and stabilize the vector for prokaryotic propagation. Lastly, the terminal non-homology Diphtheria Toxin A (DTA) chain negative selection cassette was removed, allowing for the complete targeting vector to be functionally tested via cell culture or *in utero* electroporation, which was not possible with the presence of the terminal DTA selection cassette without an additional subcloning step. Redesigned vectors and their Addgene ID numbers are outlined in Table 1 for public distribution.

**Figure 1.**
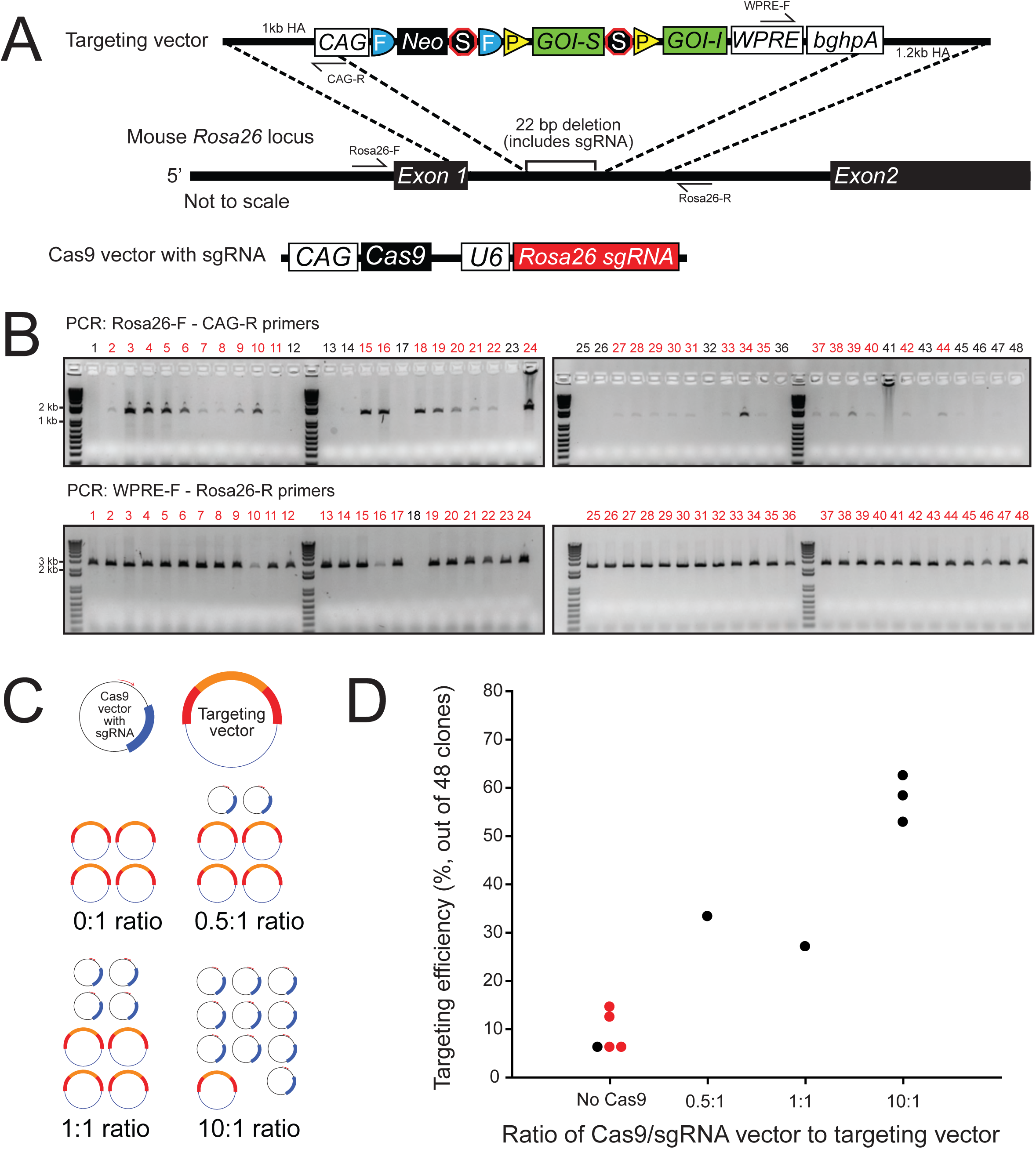
Generation of intersectional *Rosa26* mouse lines. **A)** Targeting schematic showing the modular targeting vector containing a 1kb 5’ homology arm, *CAG* promoter, *FRT*-flanked neomycin and stop cassette, *LoxP*-flanked (optional) subtractive gene of interest (GOI-S) and stop cassette, intersectional gene of interest (GOI-I), *WPRE*, *bgh poly A* element, and 1.2kb 3’ homology arm. The full intersectional cassette was knocked into the *Rosa26* locus, with a 22 bp deletion of the CRISPR sgRNA. **B)** PCR genotyping of neomycin selected ES cell clones. Targeting knock-in was determined using PCR primers that spanned from outside the *Rosa26* homology arms to either the *CAG* promoter (5’ end) or *WPRE* (3’ end). Amplification of a band indicates targeting. Shown are results from a targeting event with over 60% targeting efficiency (*RR7*). **C)** Four conditions with different Cas9:targeting vector ratios were used in our initial study: a 0:1 with no Cas9, 0.5:1, 1:1, and 10:1 **D)** Targeting efficiency results from the different ratios. The 10:1 Cas9:targeting vector ratio showed a 5-10 fold increase over an electroporation with no Cas9. Shown in red are results from previous electroporations using the traditional Rosa26 targeting vector with more commonly used longer homology arms.

**Table 1.**
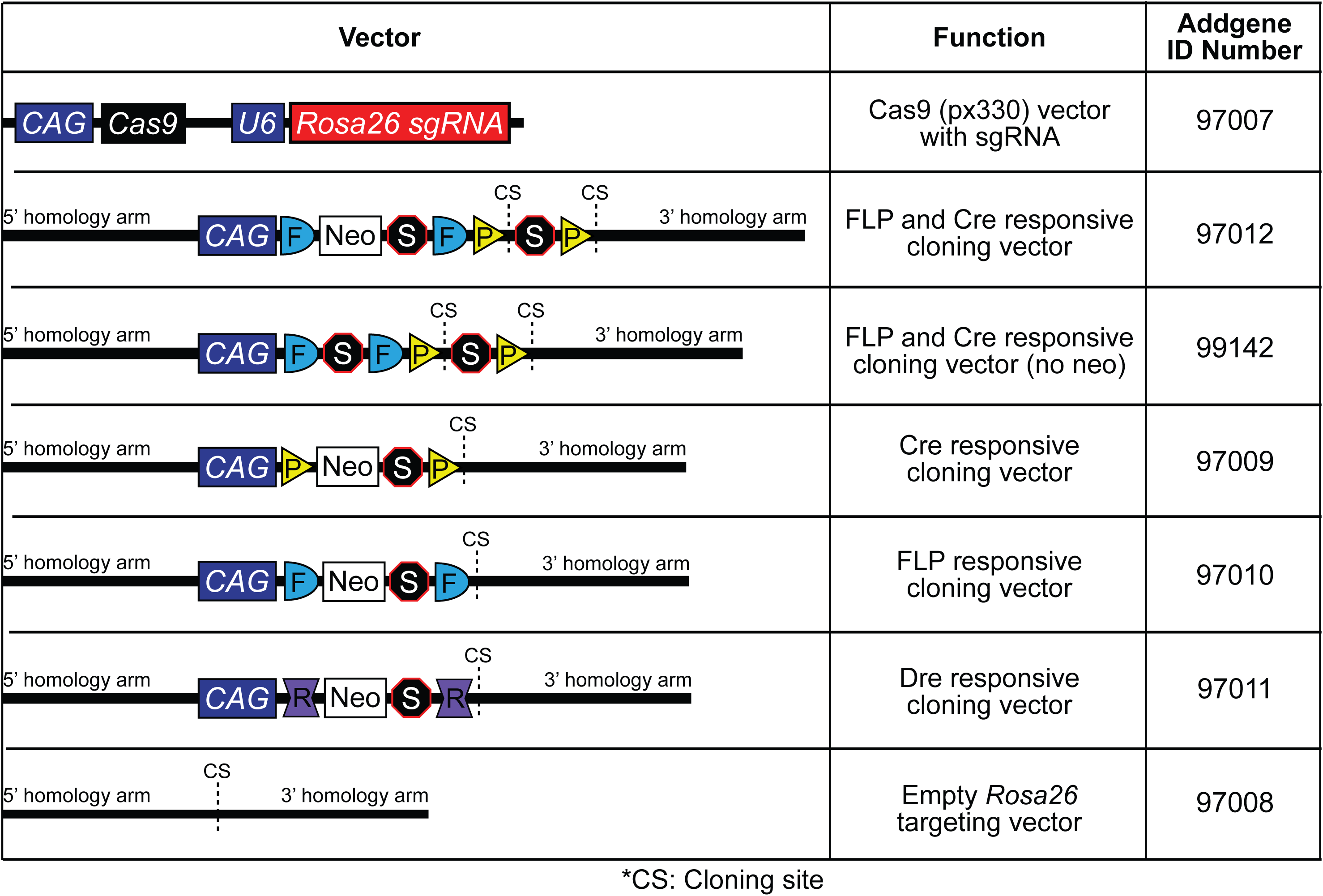
Publicly available modular targeting vectors for rapid generation of recombinase-responsive ES cells and mouse lines.

| Vector | Function | Addgene ID Number |
| --- | --- | --- |
|  | Cas9 (px330) vector with sgRNA | 97007 |
|  | FLP and Cre responsive cloning vector | 97012 |
|  | FLP and Cre responsive cloning vector (no neo) | 99142 |
|  | Cre responsive cloning vector | 97009 |
|  | FLP responsive cloning vector | 97010 |
|  | Dre responsive cloning vector | 97011 |
|  | Empty *Rosa26* targeting vector | 97008 |

\*CS: Cloning site

### Embryonic stem cell electroporation

We optimized traditional but widespread ES cell targeting by applying CRISPR/Cas9-mediated HDR (54). Our initial electroporation (EP) experiments using an earlier, more complex version of the *Rosa26* intersectional targeting vector (containing the longer 4.2 kb 3’ homology arm, DTA, and no CRISPR/Cas9) showed targeting rates ranging from 6-14% (over the course of four electroporations we saw targeting rates of 6/48, 3/48, 7/48, and 3/48 clones, mouse strains listed in **Table 2**). To determine the effect of CRISPR/Cas9 on targeting efficiency, we co-electroporated the *px330_Rosa26_sgRNA* vector expressing Cas9 and a *Rosa26* specific sgRNA with the optimized *RR5* targeting vector at four different molar ratios (0:1, 0.5:1, 1:1, and 10:1), proportionally decreasing the amount of donor vector to accommodate the increasing *px330_Rosa26_sgRNA* vector per EP (18-20µg total DNA per EP for 1×10^7^ ES cells) (**Fig 1C)**. Under the 0:1 *px330* vector:donor vector condition, we saw a 6% targeting rate similar to those seen in electroporations without Cas9 and with a much longer 3’ homology arm and negative selection, suggesting that the shortening did not have a large impact on targeting efficiency under our EP conditions. Under the 0.5:1, 1:1, and 10:1 conditions, we observed 33%, 27%, and 58% targeting, respectively, suggesting that higher ratios of the Cas9 vector resulted in increased targeting efficiency (**Fig. 1D)**.

**Table 2.**
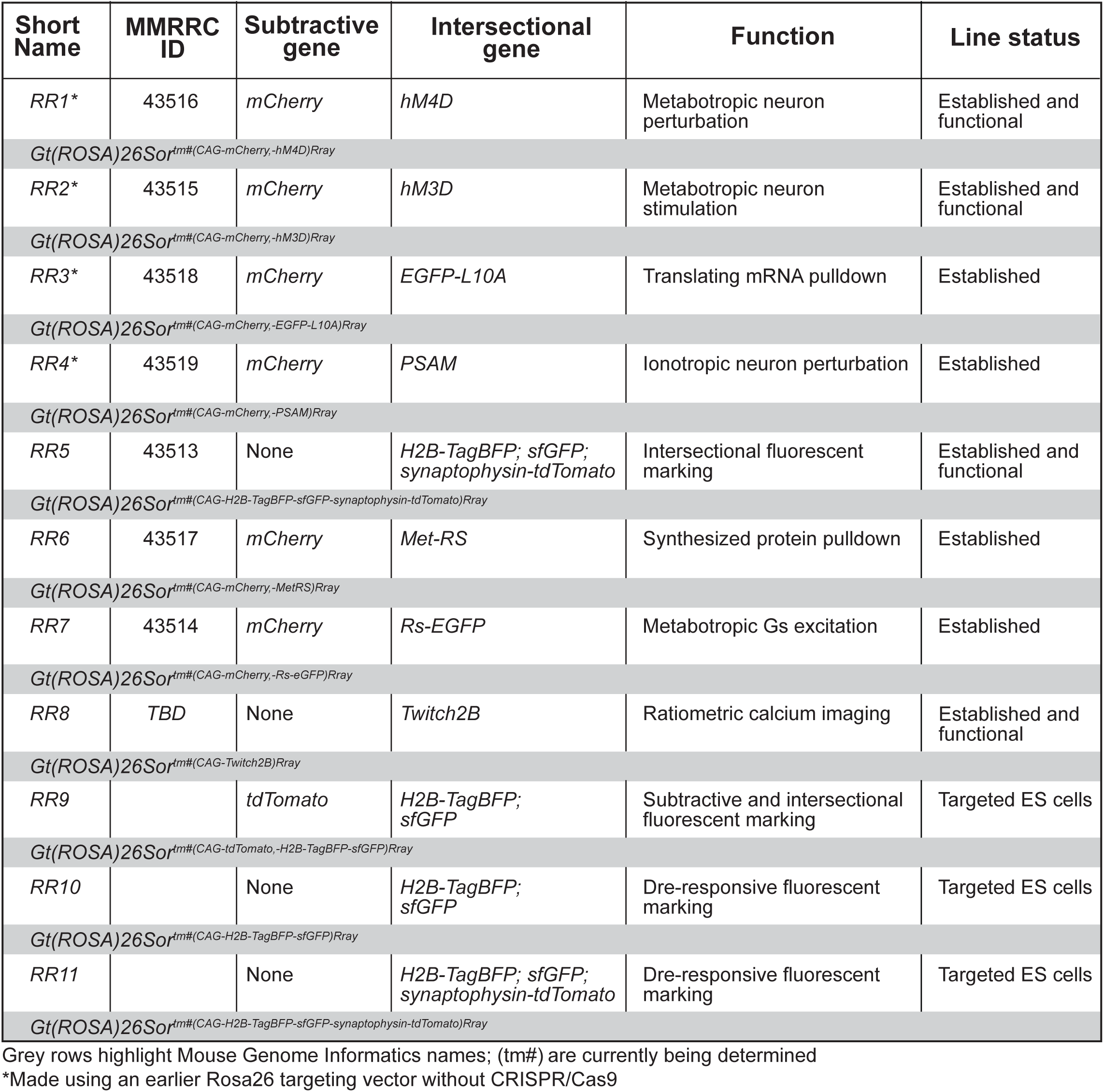
Rosa26 knock-in alleles generated with or without CRISPR/Cas9 methods. Shown are targeted Rosa26 alleles, function, length of cassette, current status, and expected MGI name (to be determined) in grey rows.

| Short Name | MMRRC ID | Subtractive gene | Intersectional gene | Function | Line status |
| --- | --- | --- | --- | --- | --- |
| RR1* | 43516 | mCherry | hM4D | Metabotropic neuron perturbation | Established and functional |
| Gt(ROSA)26Sor<sup>tm#(CAG-mCherry,-hM4D)Rray</sup> |  |  |  |  |  |
| RR2* | 43515 | mCherry | hM3D | Metabotropic neuron stimulation | Established and functional |
| Gt(ROSA)26Sor<sup>tm#(CAG-mCherry,-hM3D)Rray</sup> |  |  |  |  |  |
| RR3* | 43518 | mCherry | EGFP-L10A | Translating mRNA pulldown | Established |
| Gt(ROSA)26Sor<sup>tm#(CAG-mCherry,-EGFP-L10A)Rray</sup> |  |  |  |  |  |
| RR4* | 43519 | mCherry | PSAM | Ionotropic neuron perturbation | Established |
| Gt(ROSA)26Sor<sup>tm#(CAG-mCherry,-PSAM)Rray</sup> |  |  |  |  |  |
| RR5 | 43513 | None | H2B-TagBFP; sfGFP; synaptophysin-tdTomato | Intersectional fluorescent marking | Established and functional |
| Gt(ROSA)26Sor<sup>tm#(CAG-H2B-TagBFP-sfGFP-synaptophysin-tdTomato)Rray</sup> |  |  |  |  |  |
| RR6 | 43517 | mCherry | Met-RS | Synthesized protein pulldown | Established |
| Gt(ROSA)26Sor<sup>tm#(CAG-mCherry,-MetRS)Rray</sup> |  |  |  |  |  |
| RR7 | 43514 | mCherry | Rs-EGFP | Metabotropic Gs excitation | Established |
| Gt(ROSA)26Sor<sup>tm#(CAG-mCherry,-Rs-eGFP)Rray</sup> |  |  |  |  |  |
| RR8 | TBD | None | Twitch2B | Ratiometric calcium imaging | Established and functional |
| Gt(ROSA)26Sor<sup>tm#(CAG-Twitch2B)Rray</sup> |  |  |  |  |  |
| RR9 |  | tdTomato | H2B-TagBFP; sfGFP | Subtractive and intersectional fluorescent marking | Targeted ES cells |
| Gt(ROSA)26Sor<sup>tm#(CAG-tdTomato,-H2B-TagBFP-sfGFP)Rray</sup> |  |  |  |  |  |
| RR10 |  | None | H2B-TagBFP; sfGFP | Dre-responsive fluorescent marking | Targeted ES cells |
| Gt(ROSA)26Sor<sup>tm#(CAG-H2B-TagBFP-sfGFP)Rray</sup> |  |  |  |  |  |
| RR11 |  | None | H2B-TagBFP; sfGFP; synaptophysin-tdTomato | Dre-responsive fluorescent marking | Targeted ES cells |
| Gt(ROSA)26Sor<sup>tm#(CAG-H2B-TagBFP-sfGFP-synaptophysin-tdTomato)Rray</sup> |  |  |  |  |  |

Grey rows highlight Mouse Genome Informatics names; (tm#) are currently being determined
\*Made using an earlier Rosa26 targeting vector without CRISPR/Cas9

Since the 10:1 ratio had the highest targeting efficiency, despite having significantly less donor DNA, we used this ratio in subsequent EPs. Because the targeting efficiency was notably high, we next attempted a single electroporation with multiple vectors targeting the same locus, but containing equimolar amounts of five different cassettes, while keeping the overall *px330_Rosa26_sgRNA* vector to total donor vector ratio at 10:1. The five vectors consisted of: 1) a Cre/FLP responsive mutant methionyl-tRNA synthetase (*RR6*) for selective labeling of newly synthesized proteins; 2) a Cre/FLP responsive modified G-protein coupled receptor (*RR7*); 3) a Cre/FLP responsive bicistronic reporter with H2B-TagBFP and sfGFP separated by a p2a element, and tdTomato expressed in the subtractive population (cells that express FLP but not Cre) (*RR9*); 4) a Dre-responsive tricistronic reporter with H2B-TagBFP, sfGFP, and synaptophysin-tdTomato separated by p2a elements (*RR11*); and 5) a Dre-responsive bicistronic reporter with H2B-TagBFP and sfGFP separated by a p2a element (*RR10*). We saw a 52% targeting efficiency and successful targeting of all five cassettes at varying efficiencies (**Table 3**). Due to the lower targeting efficiency of *RR7* (2%), we attempted another 10:1 electroporation using the *RR7* donor vector alone and obtained a 63% targeting efficiency (genotyping results shown in **Fig. 1**). Thus, our results show that we are able to target and recover as many as five *Rosa* alleles in a single EP, significantly increasing efficiency and reducing costs toward intersectional mouse generation.

**Table 3.**
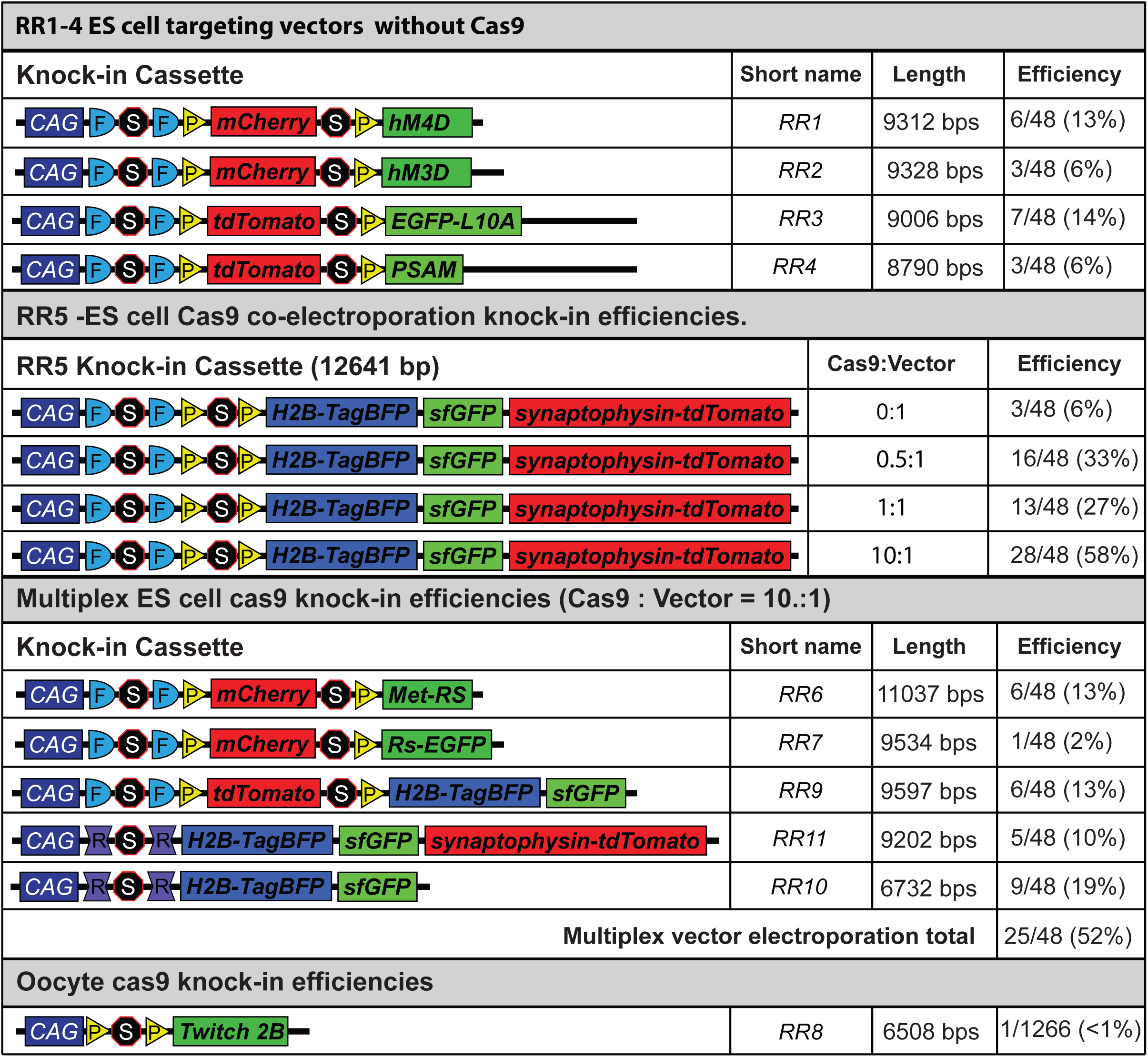
Targeting efficiencies for a multiplexed five cassette ES cell electroporation for the *Rosa26* locus.

### Oocyte targeting

Given the high efficiency of targeting, we also attempted to create a founder line (*RR8*) through direct oocyte injection of the optimized *Rosa26* targeting vector with a Cre-responsive calcium indicator, Twitch (55) (total cassette size without homology arms, 6.5kb), using pro-nuclear injection of Cas9 protein (30 ng/μl), sgRNA (20 ng/μl), and double-stranded DNA plasmid (2 ng/μl). A total of 1266 embryos were injected yielding 129 mice, of which 7 genotyped positive for the calcium indicator but only one targeted successfully (<1% targeting efficiency of mice born), suggesting that direct oocyte injection is inefficient with this system under the specified parameters.

### Off-target analysis

Genomic sequences that are similar to the sgRNA used for targeted double stranded breaks may cause unintended gene mutations or editing at off-target sites. To account for this possibility, we predicted the off-target sites for each sgRNA using the crispr.mit.edu tool and selected the top 5 sites for follow up. We PCR amplified and sequenced these loci from three correctly targeted ES cell clones and did not detect any genetic changes (**Fig. 2**). While we cannot rule out off-target effects in other loci, these data suggest that off-target effects are not prevalent in this setting.

**Figure 2.**
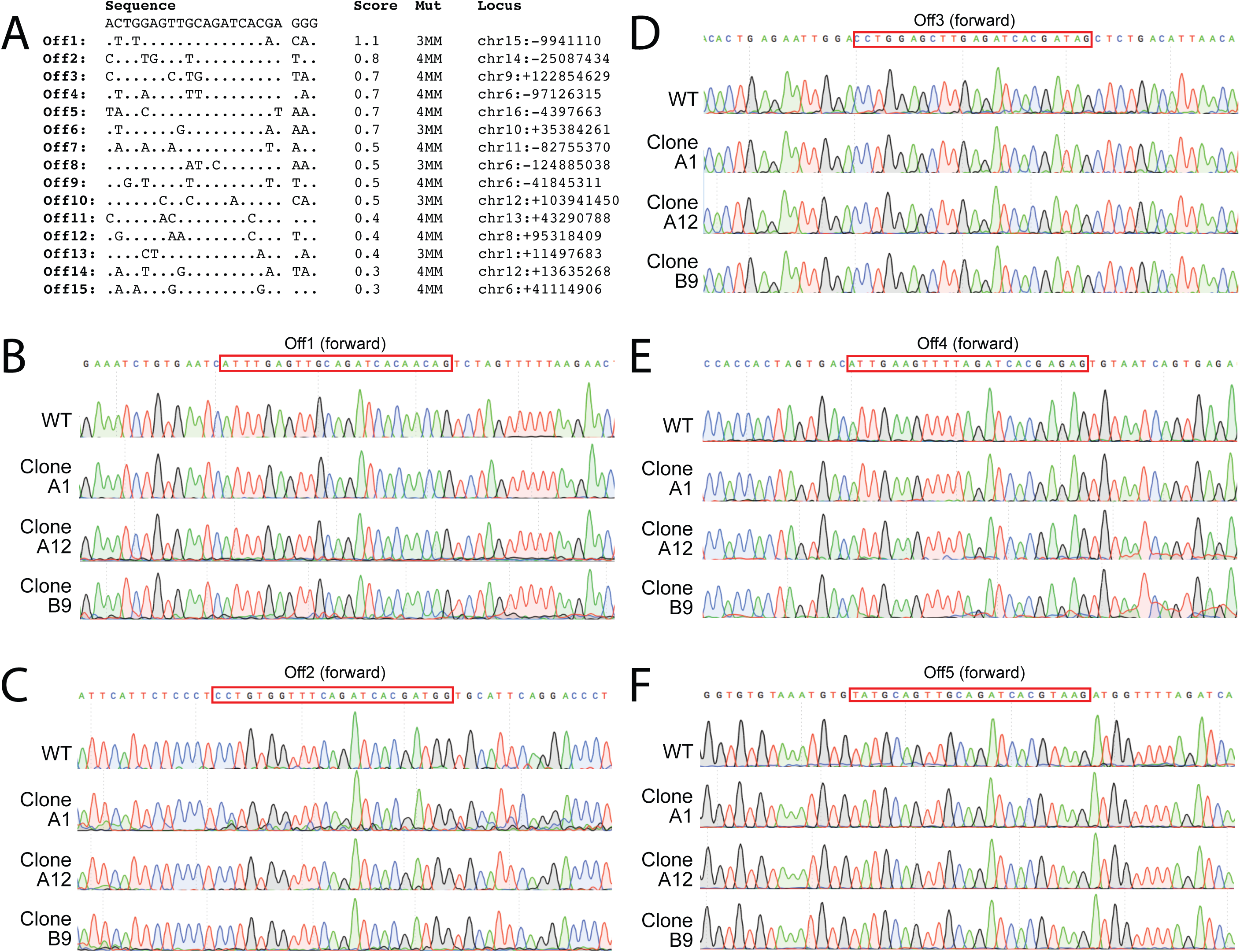
Analysis of *Rosa26* sgRNA off-target sites. No mutations were seen in the top 5 potential off-target sites. **A)** List of top 10 potential off-target sites as determined by the Optimized CRISPR tool, base pair mismatches, and location in the genome. **B-F)** Sequence chromatograms of each off-target site, showing the correct sequence for each of the 3 selected clones that were injected into blastocysts as compared to the wildtype sequence.

### Mouse line derivation

The goal of our efforts was to produce freely available genetic tools that delineate and access distinct populations for multi-faceted circuit mapping or functional characterization studies. With this toolkit in hand, we collectively produced 11 intersectional or Dre-responsive alleles available as either mouse lines (8) or ES cells (3) **(Table 2)**. To perturb neuron function, we produced three lines that express metabotropic DREADD receptors, hM4D (*RR1*), hM3D (*RR2*), and Rs-EGFP (*RR7*), and a fourth line expressing the ionotropic PSAM receptor (*RR4*) (24,56,57). Two lines enable molecular characterization: one expresses the EGFP-L10A fusion protein for ribosomal affinity purification and capture of translating mRNAs (*RR3*) (58), while the second line expresses methionyl-tRNA synthetase (Met-RS) that incorporates an artificial amino acid into nascent peptides (*RR6*) (13). One line enables ratiometric calcium imaging via expression of Twitch2B (*RR8*). Our neuro-anatomical mapping lines described below also incorporate a tagged histone (H2B), offering the possibility of chromatin affinity isolations. We created four alleles of differing Cre, FLP, and Dre recombinase responsive configurations that fluorescently mark distinct cellular compartments for unambiguous cell counts, morphological characterizations, and projection mapping (*RR5, RR9, RR10, RR11)*. The derivation of lines *RR5*, *RR6*, and *RR7* demonstrated that ES pluripotency was maintained, and that germline transmission was not diminished in our Cas9-mediated single and multiplexed electroporations after correctly targeted clones were selected and injected into blastocysts for chimera generation.

### Select mouse line characterization

Upon dual recombinase expression, the *RR5* tricistronic multi-color reporter allele expresses three spectrally separated and modified fluorescent proteins to highlight the nucleus in blue, fill the neuron in green, and emphasize pre-synaptic contacts in red. At the site of dual recombinase expression, targeted cells are brightly labelled with TagBFP, sfGFP, and tdTomato where TagBFP fluorescence is constrained to the nucleus by an H2B fusion, sfGFP is unmodified so that fluorescence is seen throughout the cell body, and tdTomato is fused to synaptophysin (59) so that fluorescence is excluded from nuclear areas and primarily seen in projection areas. Individual nuclei can be resolved using blue fluorescence and co-localization with sfGFP-labeled somata.

We used *RR5* to evaluate the functional activity and specificity *in vivo* of our CRISPR/Cas9 approach to generating mouse lines in three distinct contexts; 1) germline recombinase expression, 2) viral recombinase expression, and 3) combinatorial retrograde viral and germline recombinase expression to target single gene defined neurons by a specific projection field. First, to demonstrate genetically-restricted, germline expression of recombinases in the intersectional *RR5* line, we bred the *RR5* line to a double *Dopamine-Beta-Hydroxylase (DBH)^p2aFLPo^*(54); *Bactin-Cre* recombinase driver to express the tricistronic fluorescent cassette in *DBH-*positive noradrenergic (NA) neurons in the brainstem **(Fig. 3 A-H, Supplemental Figure 1 A-B)**. In both the locus coeruleus (**Fig. 3 A-D)** and the A5 nucleus (**Fig. 3 E-H)** as well as all other noradrenergic nuclei (not shown), we could cleanly resolve blue nuclei, green cells, and red puncta without the need for antibody enhancement. Additionally, we bred an intersection of double *Vgat-Cre; Vglut2-FLPo* recombinase expression to the *RR5* line and found the entopeduncular nucleus labeled green with local projections labeled red and distal projections in the lateral habenula labeled red (DAPI was applied to help delineate the target field, obscuring the genetic TagBFP signal)(60, 61) (**Fig. 3 I-P**).

**Figure 3.**
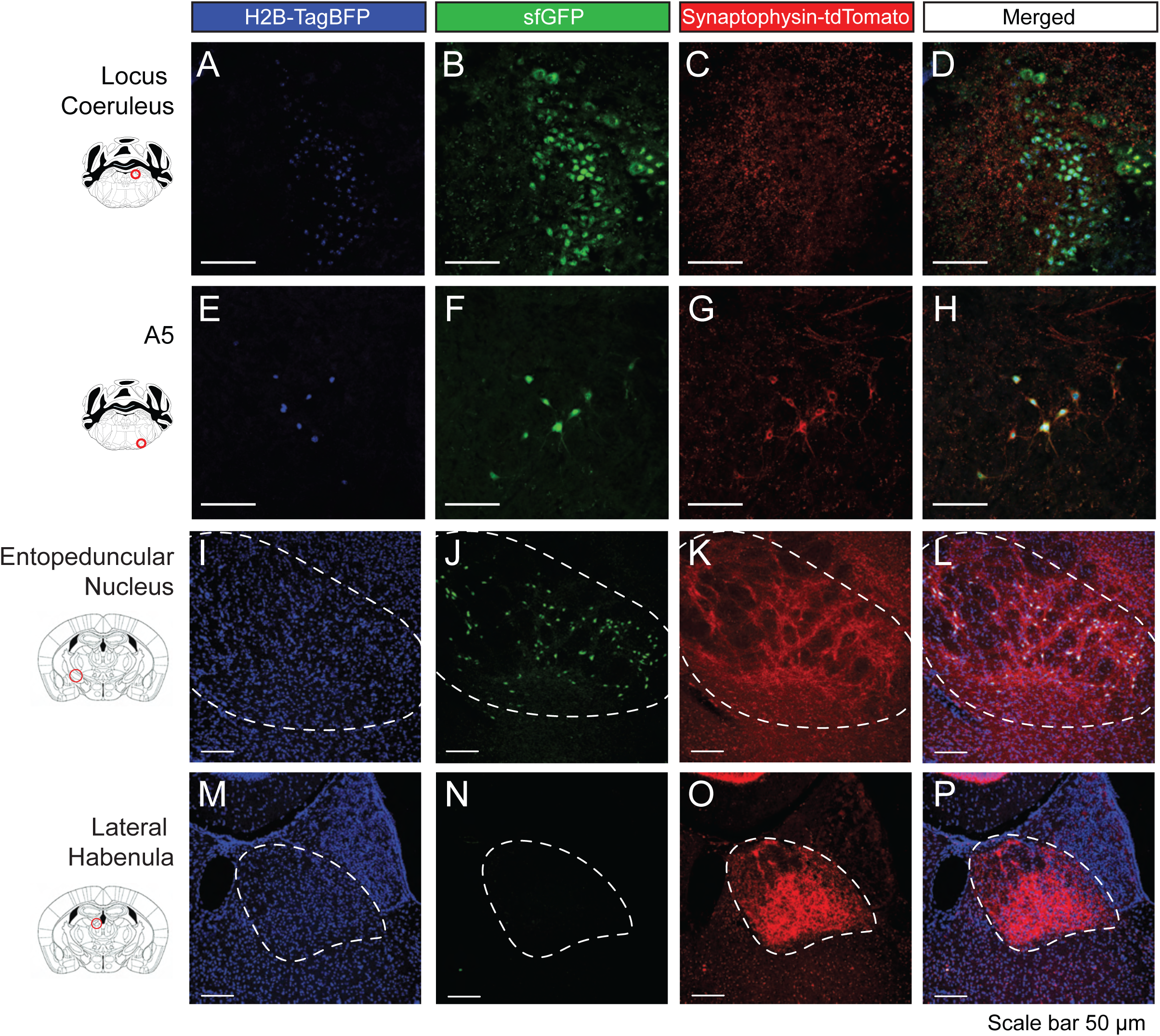
Germline Cre/FLP expression in *RR5* adult mice to fluorescently label three cellular compartments in targeted cells. **A-H)** *RR5* mice bred to *DBH^p2aFLP^; Bactin-Cre* mice express Cre ubiquitously and FLP in all noradrenergic (NA) neurons, thus marking only NA neurons, including the locus coeruleus (**A-D**) and A5 nucleus (**E-H**). **I-P)** *RR5* mice bred to *Vgat_Cre; Vglut2-FLPo* mice with dual recombinase expression imaged in the entopeduncular nucleus (**I-L**) and lateral habenula (**M-P**). In targeted cells that express both Cre and FLP, cell nuclei are labeled with TagBFP, cell soma and processes are labeled with sfGFP, and pre-synaptic contacts are labeled with tdTomato.

Second, we verified viral expression of Cre and FLP recombinase where adult *RR5* mice were stereotaxically injected with equal titer amounts of AAV viruses expressing Cre and FLP (**Fig. 4A, Supplemental Figure 1 C**) into the dentate gyrus **(Fig. 4 B-I)**, amygdala (**Fig. 4 J-Q**), and olfactory bulb (**Fig. 4 R-U**). Clear expression of all three fluorescent proteins were resolved at the site of injection to the appropriate cellular locations. Third, defining properties of the subpopulations can be extended to include any combination of gene expression or anatomical location and projection target when injection of retrograde viral vectors is applied. To utilize this option, we genetically marked a neuronal subtype defined by the partial overlap of a genetic selector and projection target by injecting canine adenovirus 2 (CAV2)-Cre virus (62, 63) into the basolateral amygdala of *DBH^p2aFLPo^; RR5* mice that express only FLP in all noradrenergic neurons. CAV2-Cre virus efficiently transduces axon terminals, thus genetically marking presynaptic neurons. In this context, only noradrenergic neurons expressing DBH that project to the amygdala express both Cre and FLP and the resulting tricistronic fluorescent cassette (**Fig. 5 A)**. We only observed recombination in the brainstem noradrenergic nuclei and in the locus coeruleus, primarily on the ipsilateral side (**Fig. 5 B-E**) with some sparse marking on the contralateral side (**Fig. 5 F-I**), in agreement with a prior study (64). Overlapping red and green puncta (with no blue) could be seen in the injected amygdala, suggesting that the marked neurons project to the amygdala, as expected given the nature of the CAV2-Cre virus (**Fig. 5 J-M).** We also observed collateral projections to several additional areas in the mid and forebrain, including the dorsal raphé (**Fig. 5 N-Q)**, reticulotegmental pontine nuclei **(Fig. 5 R-U)**, dentate gyrus **(Fig. 5 V-Y)**, and olfactory bulb (**Fig. 5 Z-CC).** This restriction by retrograde selection was also achieved in other regions of the brain with a second viral vector. *Vglut2_Cre; RR5* mice were injected with retro-AAV-Ef1a-FLPo into the lateral hypothalamus (**Fig. 6 A**) where tricistronic expression was clearly visualized in the cingulate gyrus (**Fig. 6 B-E**), piriform cortex (**Fig. 6 F-I**), and medial habenula (**Fig. 6 J-M**), indicating these regions as pre-synaptic inputs to the lateral hypothalamus. Together, these results are in agreeance with previous connectivity studies (64).

**Figure 4.**
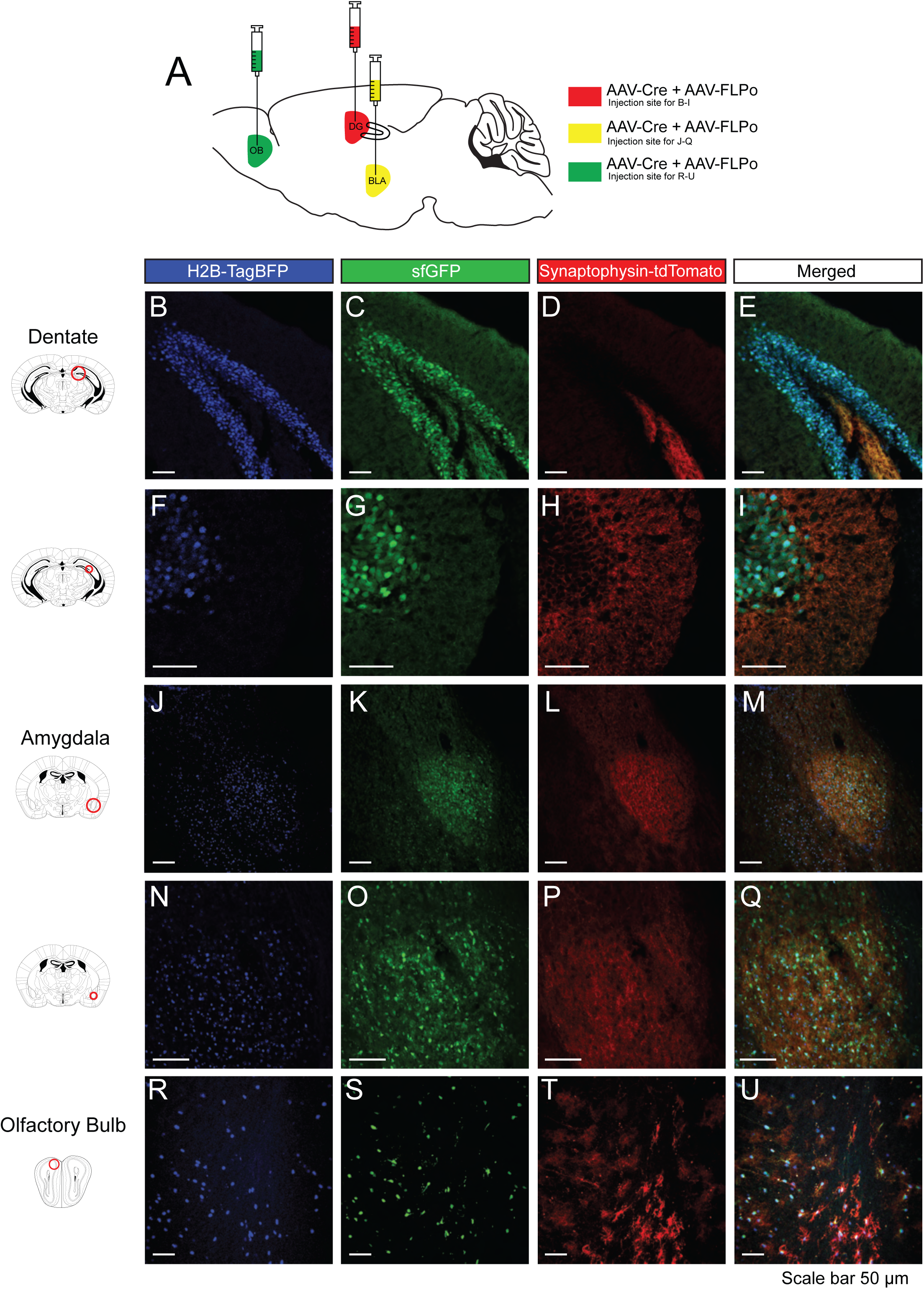
Viral-mediated expression of Cre or FLP recombinase in *RR5* adult mice to simultaneously, fluorescently label three cellular components. *RR5* mice injected with equal titers of AAV-Cre and AAV-FLP viruses into the dentate gyrus (**A-H**), basolateral amygdala (**I-P**), and olfactory bulb (**Q-T**).

**Figure 5.**
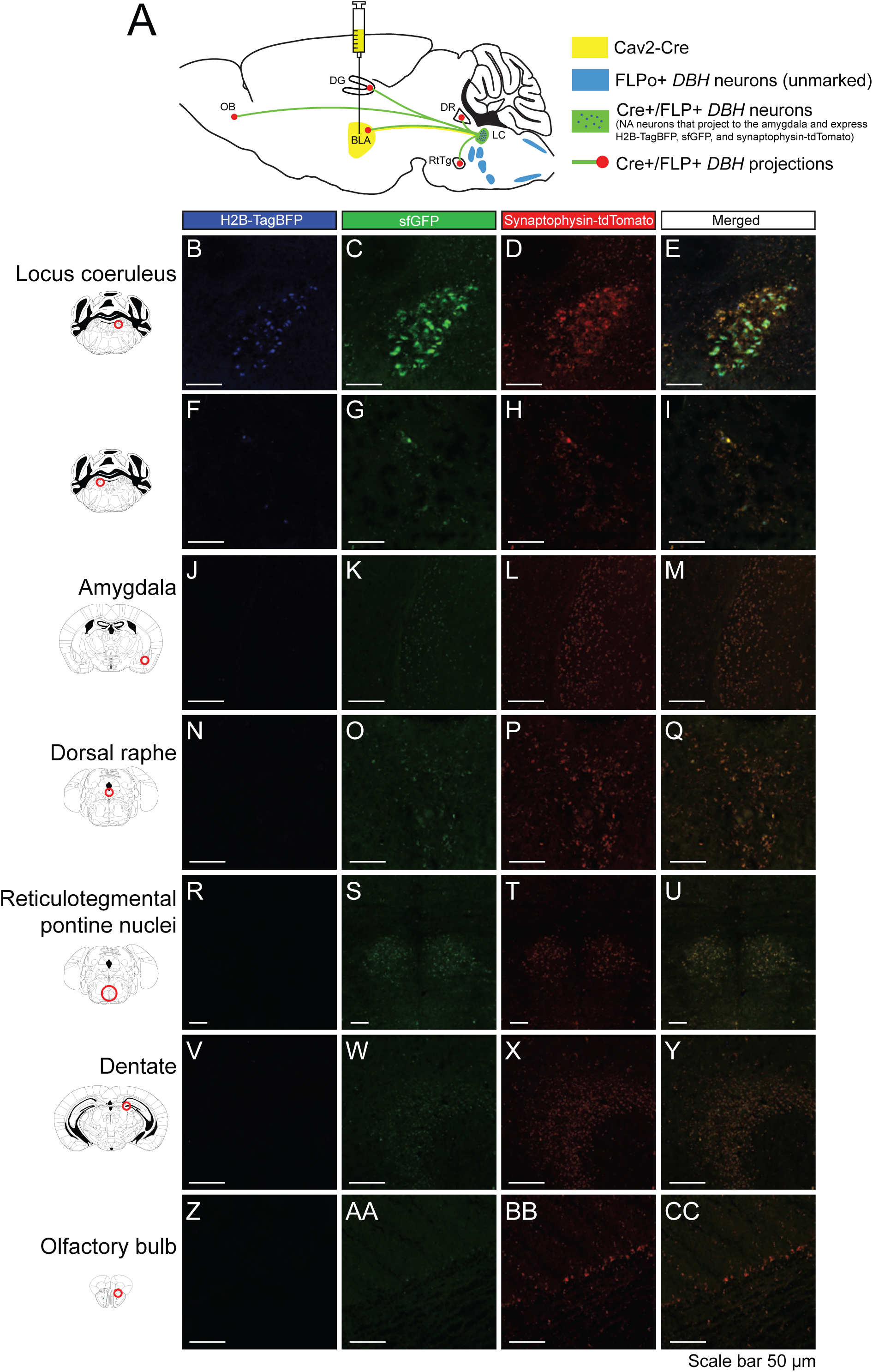
Retrograde-viral Cre and germline FLP mediated expression of dual recombinases for tricistronic FP expression on *RR5* background. **A)** *RR5; DBH^p2aFLPo^* mice were injected with CAV2-Cre virus into the basolateral amygdala. All *DBH* noradrenergic neurons express FLP, but only those noradrenergic neurons projecting to the injected amygdala will also express Cre. Marked double Cre/FLP positive cells will express H2B-TagBFP highlighting the nucleus in blue, sfGFP filling the cell including the axon, and synaptophysin-tdTomato labeling presynaptic contacts. BLA: Basolateral amygdala. DG: Dentate gyrus. DR: Dorsal raphé. LC: Locus coeruleus. OB: Olfactory bulb. RtTg: Reticulotegmental pontine nucleus. *DBH* positive neurons that project to the amygdala arise primarily from the ipsilateral locus coeruleus (**B-E**) with some sparse labeling in the contralateral locus coeruleus (**F-I**). Red puncta overlapping with green but lacking blue marked nuclei (indicative of projections from the marked population) can be seen in a variety of mid- and forebrain areas, including the injected amygdala (**J-M**), the raphé nucleus (**N-Q**), reticulotegmental pontine nuclei (**R-U**), dentate gyrus (**V-Y**), and olfactory bulb (**Z-CC**).

**Figure 6.**
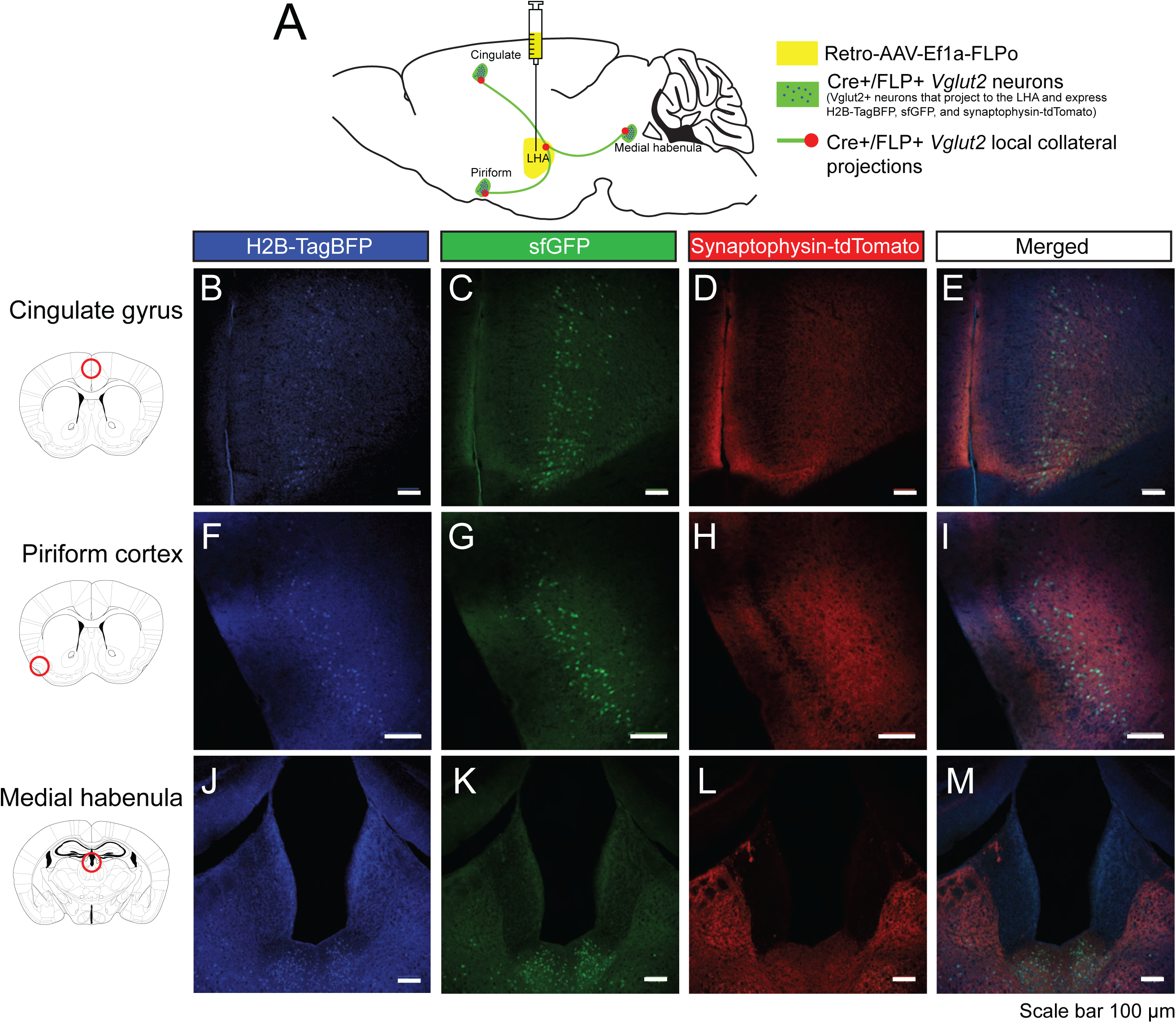
Retrograde-viral FLP and germline Cre mediated expression of dual recombinases for tricistronic FP expression on *RR5* background. **A)** *RR5; Vglut2^Cre^* mice were injected with retro-AAV-Ef1a-FLPo virus into the lateral hypothalamus. All *Vglut2* neurons express Cre, but only those Vglut2 neurons projecting to the injected LHA will also express FLP. Marked double Cre/FLP positive cells will express H2B-TagBFP highlighting the nucleus in blue, sfGFP filling the cell including the axon, and synaptophysin-tdTomato labeling presynaptic contacts. LHA: Lateral hypothalamic area. Vglut2 positive neurons that project to the LHA arise from the cingulate gyrus (**B-E**), piriform cortex (**F-I**), and medial habenula (**J-M**).

Before *in vivo* characterization of the DREADD systems coded by *RR1* and *RR2*, we first examined the ability of the *RR1 and RR2* lines to modulate dopamine beta-hydroxylase (DBH)- defined noradrenergic neuron activity at the cellular level. We conducted whole-cell recordings from P30-P60 locus coeruleus neurons expressing one of 3 DREADD receptors: hM3D (RR2), hM4D (RR1), or Di (previously published). Each cassette exists with FRT and LoxP bound stop cassettes, so each line was initially bred with Bactin_FLPe to remove the FLP-dependent stop cassette (65). The RC::FrePe was also bred with Bactin_FLPe to remove the FLP-dependent stop cassette and then bred to the cre-only dependent DREADD lines (66). This compound allowed for fluorescent labelling of neurons where Cre recombinase, and thus DREADD receptors, are present. These compounds resulted in mice wherein GFP and a DREADD were expressed in *TgDBH-Cre-*defined cells for acute brain slice visualization (See Supplemental Figure 2 for further genetic details). *RC::PDi* (52) was included as it represents a similarly constructed hM4D intersectional allele, but lacks the modifications and optimizations made in our targeting system (**Fig. 1 A**). For example*, RC::PDi* contains an additional PGK Promoter and pA signal sequence flanking neomycin in the first stop cassette and lacks the WPRE element and pA found in *RR1* and *RR2*. We hypothesized that the addition of a Woodchuck Hepatitis Virus (WHV) Posttranscriptional Regulatory Element (WPRE) would enhance DREADD effectiveness. After bath application of 10 μM CNO, for *RR2P (hM3D); RC::epe* we observed a depolarization of membrane potentials of LC neurons (pre-CNO: -62.26±1.53, post-CNO: -55.09±2.16 mV, paired t-test: p=0.0034, n=19 neurons across 3 mice). After bath application of 10 μM CNO, for *RR1P (hM4D); RC::epe* we observed a hyperpolarization of membrane potentials of LC neurons (pre-CNO: -56.98±3.50 mV, post-CNO: -65.3137±3.60 mV, paired t-test: p=0.046, n-10 neurons across 3 mice). After bath application of 10 μM CNO, for *RC::PDi (Di); RC::epe* we observed a hyperpolarization of membrane potentials (pre-CNO: -53.71±2.93 mV, post-CNO: -56.72±3.42 mV, paired t-test: p = 0.027, n=10 neurons across 3 mice) (**Fig. 7 A-C**). These data confirmed functional expression of each DREADD system.

**Figure 7.**
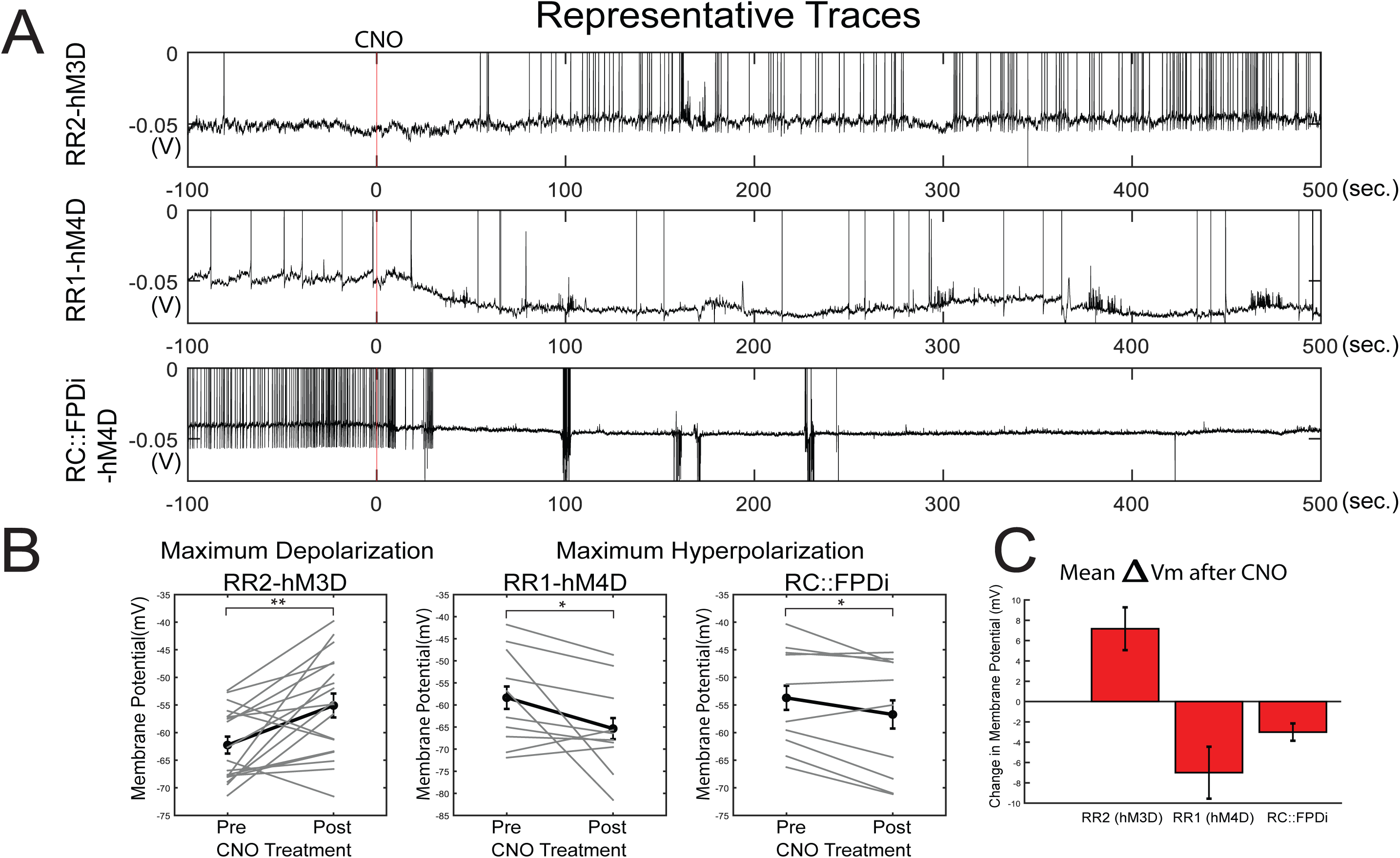
Electrophysiological characterization of CNO-DREADD mediated responses in noradrenergic locus coeruleus (LC) neurons from lines *RR1(P), RR2(P),* and *RC::FPDi (P)*. **(A)** Representative wash-on responses of LC neurons from *RR1(P) (n=19 neurons), RR2(P) (n=10 neurons),* and *RC::FPDi*(P) *(n=10 neurons)* lines in response to DREADD agonist CNO. **(B)** Membrane potentials of each LC neurons (pre-CNO vs post-CNO values). **(C)** Summary plots of absolute voltage change for each DREADD line tested following CNO treatment. (*p<0.05; **p<0.01).

We next performed whole-body plethysmography to assess changes in respiratory function caused by CNO-DREADD mediated perturbation of targeted neurons *in vivo*. Significant changes in any of the calculated respiratory variables demonstrates a measurable effect of DBH-defined noradrenergic neuronal activity on respiratory control and/ or ventilatory response to CO_2_. If change was seen in room air, then basal respiratory control was affected by the change in activity of these neurons. If change were seen in 5% CO_2_, then changes in the activity of these neurons affected the hypercapnic ventilatory response. *DBH-Cre* defined NA neurons were inhibited by crossing mouse line *RR1P* to *Tg(Dbh-cre)KH212Gsat (TgDBH-Cre)* mice. We observed no changes under room air conditions (**Fig. 8 A)**. However, under hypercapnic conditions, we saw a reduction in ventilation (**Fig. 8 B)**, with decreases in V_T_ (Tidal volume) (-26.6%, p=0.0021, **Fig. 8 D**), 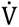_E_ (Minute ventilation) (-26.8%, p=0.026, **Fig. 8 E**), and 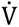_E_/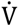_O2_ (Ventilatory equivalents for oxygen) (-28.7%, p<0.0001, **Fig. 8 G**). No significant differences were observed in body temperature between experimental animals and sibling controls (**Fig. 8 H**). Notably, sibling controls showed no difference in respiratory parameters pre- and post-CNO administration.

**Figure 8.**
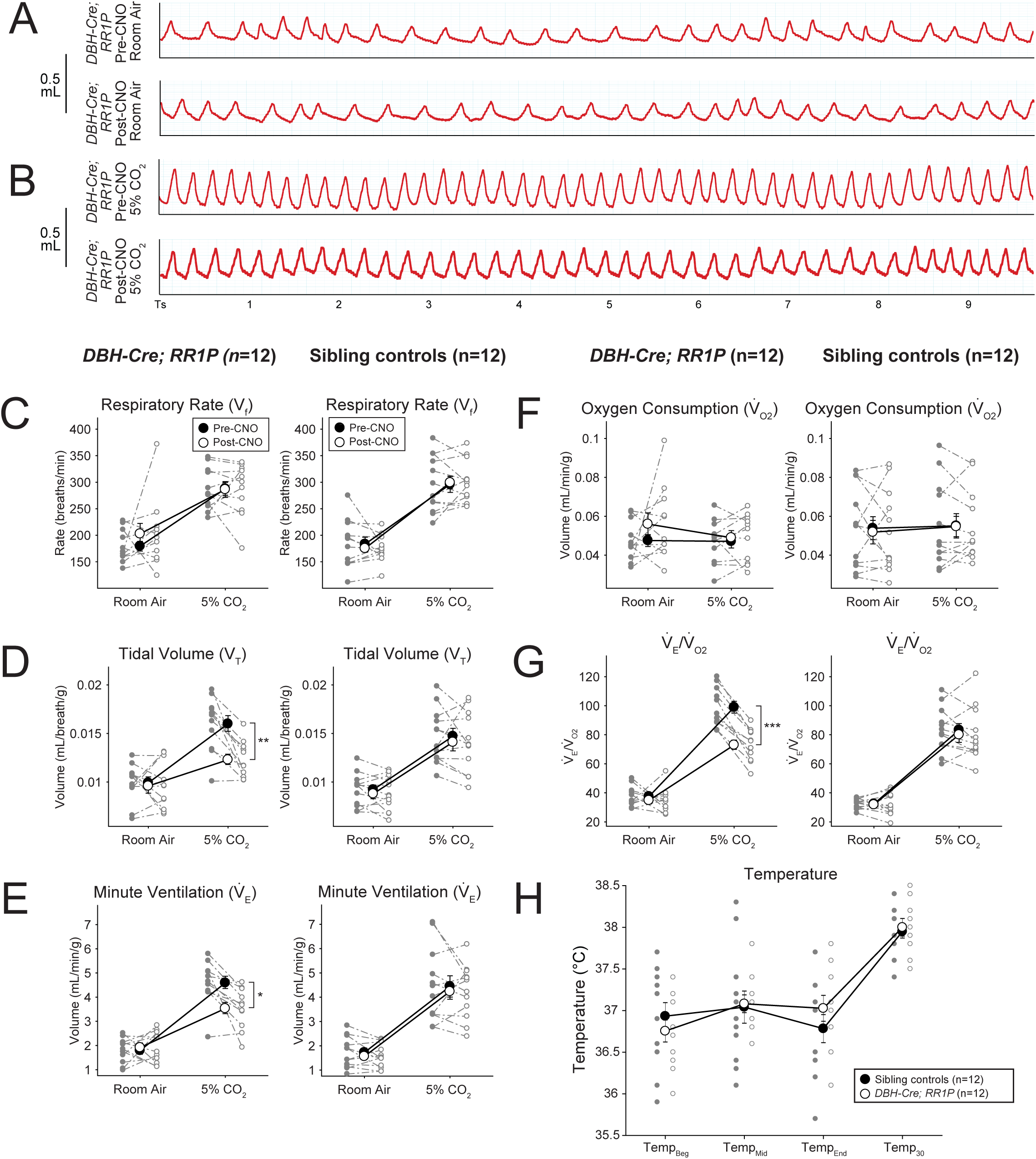
CNO-hM4D mediated perturbation of noradrenergic *DBH-Cre* neurons results in a reduced hypercapnic response. **A)** Representative breathing traces from a *DBH-Cre; RR1P* animal before and after CNO administration under hypercapnic conditions (5%CO_2_/21%O_2_/74%N_2_). **B-F)** Quantification of respiratory and metabolic parameters under room air and hypercapnic conditions in *DBH-Cre; RR1P* animals and sibling controls. Measured values include respiratory rate (**B)**, tidal volume (**C)**, minute ventilation **(D)**, oxygen consumption **(E)**, and minute ventilation normalized to oxygen consumption **(G)**. No difference was seen under room air conditions but *DBH-Cre; RR1P* animals showed a deficit in volume, minute ventilation, and minute ventilation normalized to oxygen consumption under hypercapnic conditions. **G)** No difference in temperature was seen between *DBH-Cre; RR1P* animals and sibling controls. *p<0.05; **p<0.01; ***p<0.001.

We then tested the applicability of the *RR2P* line *in vivo.* After CNO-DREADD mediated stimulation of *DBH-Cre* defined neurons, we saw significant increases under room air ventilation (**Fig. 9 A)** in V_f_ (Ventilatory frequency) (+28.1%, p=0.0076, **Fig. 9 C**), V_T_ (+43.3%, p=0.00077, **Fig. 9 D**), 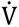_E_ (+83.6%, p<0.0001, **Fig. 9 E**), and 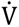_O2_ (Oxygen consumption) (+50.9%, p<0.0001, **Fig. 9 F**). The increase in 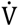_E_ was proportional to the increase in 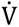_O2_, however, so 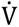_E_/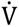_O2_ did not change **(Fig. 9 G)**. Under hypercapnic conditions, as compared to pre-CNO values, 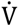_O2_ was increased (+37.2%, p=0.0015, **Fig. 9 F)**, resulting in a reduced 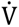_E_/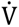_O2_ (-24.2%, p=0.04, **Fig. 9 G)**. We also observed a deficit in body temperature in *DBH-Cre; RR2P* animals 30 minutes after the end of the assay at room temperature (∼1.5 hrs after CNO injection) as compared to sibling controls (**Fig. 9 H)**, but not immediately at exit from the respiratory chamber, which is kept at a thermoneutral temperature, 30-32°C. Sibling controls showed no difference in respiratory parameters pre- or post-CNO administration. The respiratory phenotypes seen here clearly demonstrate an acute and cell autonomous involvement of the whole NA system in CO_2_ chemosensitivity and baseline metabolism while avoiding the confounds of developmental compensatory events or off-target effects.

**Figure 9.**
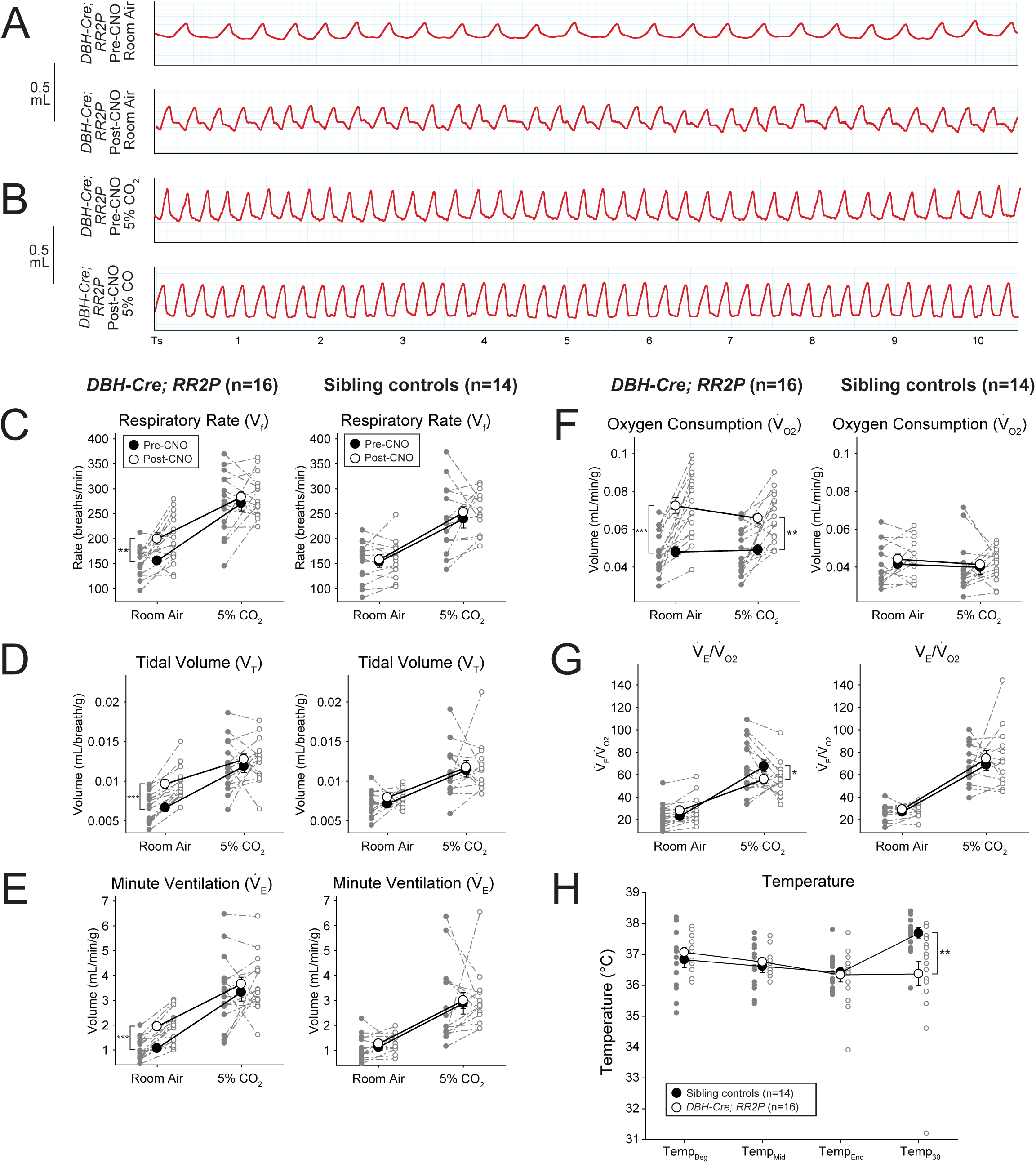
CNO-hM3D mediated perturbation of noradrenergic *DBH-Cre* neurons results in enhanced room air ventilation and a reduced hypercapnic response. **A)** Representative breathing traces from a *DBH-Cre; RR2P* animal before and after CNO administration under room air conditions (21%O_2_/79%N_2_). **B)** Representative breathing traces from a *DBH-Cre; RR2P* animal before and after CNO administration under hypercapnic conditions (5%CO_2_/21%O_2_/74%N_2_). **C-G)** Quantification of respiratory and metabolic parameters under room air and hypercapnic conditions in *DBH-Cre; RR2P* animals and sibling controls. Measured values include respiratory rate **(C)**, tidal volume **(D)**, minute ventilation **(E)**, oxygen consumption **(F)**, and minute ventilation normalized to oxygen consumption **(G).** After CNO administration, *DBH-Cre; RR2P* animals showed increased rate, volume, minute ventilation, and oxygen consumption under room air conditions. Under hypercapnic conditions, *DBH-Cre; RR2P* animals showed increased oxygen consumption resulting in a reduced minute ventilation to oxygen consumption value. **I)** *DBH-Cre; RR2P* animals showed a significant deficit in temperature 30 minutes after the end of the assay. *p<0.05; **p<0.01; ***p<0.001.

Finally, we characterized a fourth line, produced by oocyte CRISPR-mediated recombination to express the Twitch2B ratiometric calcium indicator upon Cre recombinase expression. Calcium signaling was recorded in outer granulosa cells surrounding an oocyte taken from a germline recombined (*RR8; Bact_Cre*) mouse line (**Fig. 10 A**). Upon application of 10µM ionomycin in the presence of 10mM CaCl_2_, the YFP/CFP ratio increased more than 3-fold (**Fig. 10 B**). The data demonstrate how the targeting system here can be used to build conditional mouse lines that are useful beyond the nervous system and throughout the body (67).

**Figure 10.**
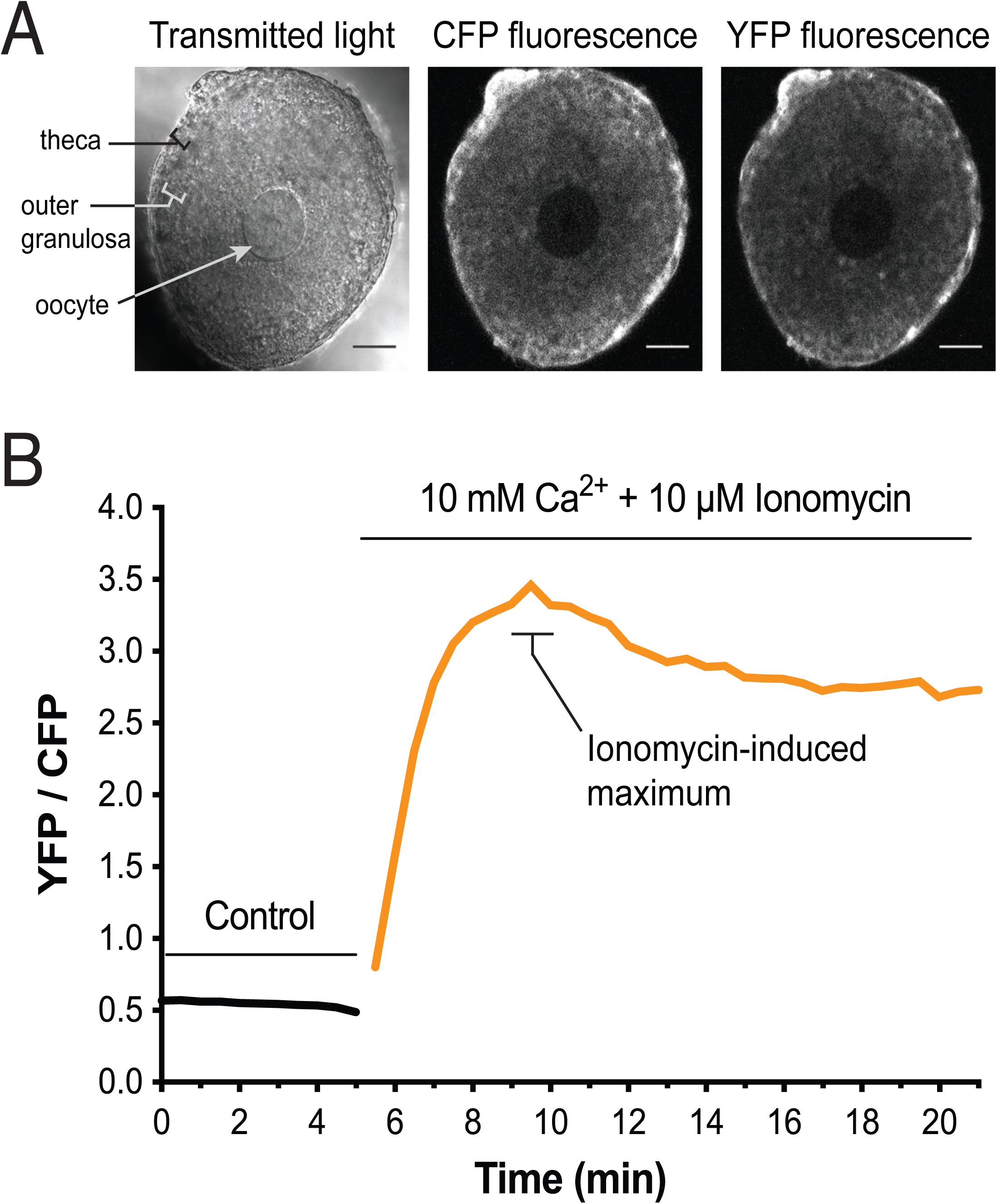
Expression and function of Twitch-2B in the granulosa cells of mouse ovarian follicles. (**A**) A follicle from a mouse expressing Twitch-2B (homozygote). A follicle consists of multiple layers of granulosa cells surrounding an oocyte in the center. Twitch-2B is uniformly expressed in the granulosa cells but is not detectable in the oocyte. A layer of theca cells, adhering to the outside of the follicle, expresses Twitch-2B at a higher level. Scale bar = 50 µm. **(B)** ∼6-fold increase in YFP/CFP ratio in the outer granulosa cells after application of 10 µM ionomycin in the presence of 10 mM CaCl_2_. Representative of 4 follicles.

Collectively, these data highlight how our single recombinase and intersectional lines can be used to functionally assess dispersed and difficult to access populations, setting the stage for further characterization and dissection of a variety of systems in a given measured outcome through intersectional subdivision by genetic or viral methods.

## DISCUSSION

### Vector design and optimization

Our tricistronic mouse line vector (*RR5*) was a complex vector to design. Thus, the discussion of its construction applies to the other simpler vectors and best describes choices made to optimize the vectors. Our initial goal was to create an intersectional mouse strain expressing three fluorescent components to highlight the nucleus, whole neuron, and presynaptic contacts, enabling facile study of cell counts, cellular morphology, and axonal and projection targets (*RR5* consisting of H2B-TagBFP, sfGFP, and synaptophysin-tdTomato separated by p2a elements) without the need for signal amplification using antibodies. However, likely due to the high number of repetitive elements in the *Rosa26* 3’ homology arm and the high complexity and repetitive components of the intersectional cassette (i.e. two stop cassettes consisting of multiple SV40 polyadenylation sequences, tdTomato protein dimer, double loxP and FRT sites, and two p2a self-cleaving peptides separating the elements), we had no success in stably subcloning the intersectional cassette into a low copy vector (p15a ori) containing the original *Rosa26* homology arms (1.1kb 5’ homology arm and 4.2 kb 3’ homology arm) and DTA negative selection gene (52). Thus, we sought to make several improvements on the prior approaches used to build intersectional genetic mice. First, we sought to stabilize the vector by shortening the homology arms and eliminating repetitive genomic elements found at the *Rosa26* locus. Second, we removed the DTA negative selection terminal non-homology to eliminate the need to validate intersectional cassettes separately before placing the cassette in the targeting vector. Third, we reduced the size of the selection cassette. Prior intersectional constructs carried an additional PGK promoter and BGHpA tail for neomycin expression. We eliminated both promoter and BGHpA as the functions of these elements can easily be served by the CAG promoter and SV40pA in the stop cassette. With smaller homology arms, it was possible that our targeting efficiency would fall. Therefore, we introduced a CRISPR/Cas9 targeting strategy to enhance homologous recombination in either ES cells or oocytes. Surprisingly, in ES cells, the shorter homology arms performed equally well as the full-length homology arms (see discussion below). The resulting strategy offered a simplified, smaller, and highly stable vector that remains efficient at targeting the *Rosa26* locus. Through this optimization process we built a set of baseline vectors that can be used to build any intersectional genetic targeting allele rapidly and inexpensively by simply cloning in a cDNA of interest into one or both of the multiple cloning sites for both intersectional and subtractive expression.

### Embryonic stem cell electroporation

While CRISPR strategies are widespread for knockout mutations and small deletions and insertions in mouse zygotes, consistent knock-in of large targeting cassettes (>4kb) still remains a challenge with fewer but increasing successes (68–70). To address this limitation, our new multiplexed methodology enabled us to: **1)** simplify our base targeting vector to less than 5kb in total length; **2)** see successful targeting of added cassettes up to 11kb; **3)** increase the rate of targeting by 5-10 fold over previous *Rosa26* targeting attempts (that used a significantly longer 3’ homology arm) under our ES cell strain and conditions; and **4)** further reduce cost by co-electroporating five different targeting vectors in a single ES cell electroporation that were easily resolved to produce new mouse lines. While targeting rates for the *Rosa26* locus can vary greatly in the literature, we compare only to our own experiments, as significant variability in targeting efficiency from lab to lab or facility to facility can arise from the several factors in ES cell electroporation ranging from electroporation conditions (e.g. buffer ionic strength, adjuvants, field strength, field duration, field shape, square wave vs exponential decay, etc.) to culture conditions (ES cell strain and genetic background, feeder type, media composition, inclusion of LIF, etc.). Notably, when we co-electroporated a CRISPR**/**Cas9 plasmid to facilitate targeting, there was a proportional increase in targeting efficiency despite reduced amounts of targeting vector (total DNA in the EP was capped at 18-20 µg per EP). At the highest efficiencies, we could further subdivide the small amount of targeting vector across multiple plasmids, which allowed us to introduce and recover as many as five vectors in a single EP. This multiplexed approach significantly reduced the costs of mouse production via ES cells. With these goals met, multiplexed ES cell targeting enables the rapid and high throughput production of intersectional alleles that can be readily distributed throughout the mouse research community for further studies.

### Oocyte targeting

Oocyte injection was only attempted with one line, *RR8*. Less than 1% of the embryos contained a properly targeted vector, which in our experience suggests that direct oocyte injection is not efficient. Future optimization and the development of new technologies may increase the efficiency of targeting for more rapid generation of transgenic mice; for example, a study showed 10-20% targeting of 8-11kb cassettes to the *Rosa26* locus in oocytes using longer homology arms, while another showed successful targeting of a 5.5kb cassette into the *Rosa26* locus in rats with co-injection of two ssDNA oligos (70, 71). As noted above, there can be significant variability in targeting across labs based upon a variety of factors including background strain (vs strain from which the homology arms are derived), the use of Cas9 mRNA or protein, and site of injection (pro-nucleus vs cytoplasm, single vs 2-cell stage). Thus, our outcomes here may reflect our core facility capabilities as much as inherent limitations in consistent Cas9 mediated oocyte targeting of large constructs. The successful outcomes here and in a prior publication demonstrate the use of the system to develop lines capable of functional imaging (67).

### Select mouse line characterization

Intersectional expression of fluorescent proteins (FPs) has been used to great effect for anatomical characterizations of targeted neuron populations (66, 72). Characterizations aimed at counting or highlighting single cells or cell populations made with a single FP can be made difficult, however, in cases where cell, axon, or dendrite density (neuropil) is high. Usually, one or two FPs are deployed to either fill the cell or highlight specific features, which necessitates the use of multiple reporter alleles to clearly observe cell number, morphology, and projections. To ameliorate these issues, we sought to create a mouse line (*RR5*) that highlights three specific features in an intersectionally defined neuron. Overall, this novel three-color mouse line enables intersectional and simultaneous labeling of three subcellular compartments. The polycistronic fluorescent labeling cassette gives unambiguous cell counts and clear visualization of cell morphology, axonal projections, and synaptic contacts. The strength of expression of all three colors is likely due to the use of p2a elements, rather than IRES elements, which cause expression level reduction. Reduced expression from an IRES may stem from cryptic splice elements in the IRES that could interact with the intron in the CAG promoter or splice acceptors in some stop cassettes (73). Our use of P2A appears to bypass those issues as all three fluorescent proteins could be readily visualized without additional enhancement as compared to other approaches (38).

We also show that combinations of germline, viral, and retrograde viral expression of recombinases in the *RR5* line can all be used to clearly map the shared innervation of multiple regions by a small subset of genetic, projection, and/ or intersectionally defined neurons. In contrast to prior intersectional studies using two genetic delimiters to define a subpopulation with a single fluorescent protein (with or without an inert retrograde marker injected into separate sites in the brain), our approach enables facile collateral mapping across the whole nervous system with higher resolution that is growing in popularity (36,64,72). These results demonstrate that despite the high complexity and repetitive nature of the *RR5* cassette, we could efficiently target this allele, the allele remains stable through germline transmission, and it is responsive to Cre and FLP recombinase expression. The restriction of expression to anatomically defined NA neurons as a whole or as a projection subset further establishes the recombinase specificity of our targeting schematic. This approach should also translate equally well to further anatomical, functional, and molecular characterizations of NA and other subtypes defined by their projection patterns with the use of the additional lines reported here.

The outcomes of either inhibiting (*RR1P; TgDBH_Cre*) or stimulating (*RR2P; TgDBH_Cre*) NA neurons as defined by *TgDBH_Cre* strongly support the functionality of these two lines in an *in vivo* setting. Noradrenergic (NA) neurons are strongly implicated in control of respiratory homeostasis and chemosensitivity (8,74–80). Previous studies focused on demonstrating the role of NA neurons in breathing bear methodological caveats such as lack of resolution due to overly broad lesions or injections (81), developmental and non-cell-autonomous compensation after gene mutations, restraint or anesthesia *in vivo*, and they typically lack concurrent metabolic measurements (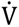_E_/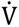_O2_) (82), thus direct comparisons are difficult. Nonetheless, previous work has been reiterated in many ways with our results here in that focal pharmacological NA lesions decreases respiratory frequency and hypercapnic ventilatory response (76). Here we show in conscious, unrestrained mice that inhibition of *TgDBH_Cre*-defined noradrenergic neurons led to significantly reduced tidal volume, minute ventilation, and 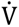_E_/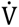_O2_ following hypercapnic exposure. On the other hand, activation of *TgDBH_Cre*-defined noradrenergic neurons led to significantly increased respiratory rate, tidal volume, minute ventilation, 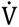_O2_, and 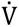_E_/ 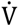_O2_ in the room air breathing condition. These changes likely stem from a combined increase in central and peripheral (sympathetic) outflow increasing overall metabolic rate (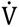_O2_). Although breathing and 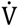_O2_ are both increased, a small but significant mismatch occurs in hypercapnia suggesting that global excitation may destabilize hypercapnic reflexes as well. Notably, after removal from the thermo-neutral recording chamber, the body temperature dropped significantly. This drop is consistent with prior studies that show central NA outflow to the hypothalamus negatively regulates body temperature (83, 84). The magnitude of changes seen for both stimulation and inhibition of *TgDBH_Cre* defined NA neurons in various breathing parameters, particularly for hypercapnia, is in line with expectations of a highly redundant chemosensory system in the respiratory network along with the fact that the NA system is neuromodulatory in nature (37,52,85–88). Altogether, the numerous breathing, metabolic, and temperature changes observed support the functionality of these lines for *in-vivo* chemogenetic manipulations. These data represent modulation of the noradrenergic system that, methodologically, significantly deviates from previously published studies interrogating this system and provides an alternative to those investigating control of breathing that reduces the influence of unintended effects.

In conclusion, the methods presented here represent a significant improvement in feasibility and affordability for labs to generate their own intersectional mouse models for application in a variety of fields. Our vector design and application allow for large cassettes to be targeted to the *Rosa26* locus with high throughput, increased efficiency, germline transmission, and preserved pluripotency. These protocols and techniques can be used to investigate most, if not all, neural circuits in the context of development and/ or disease. Implementing these techniques also serves as an alternative to cost-prohibitive commercially available vectors and mouse lines. Together, we present a facile, cost-effective method for producing gene specific intersectional mouse lines.

## CONCLUSIONS

During development, many distinct cellular subtypes arise and intercalate to create the complex cells, organs, and organ systems that constitute our behavior and physiology. Key to understanding this organization and how it may be perturbed in diseases is the ability to identify and access discrete cell subtypes in the developing and adult mouse for multi-faceted studies. It is now clear that even within narrowly defined cell types once thought as homogeneous, significant diversity is found at multiple levels including genetic and molecular signatures, activity patterns, synaptic connectivity, and projection patterns. Given this complexity, it is often difficult to define and access specific cellular populations for study, particularly during embryogenesis, where *in utero* development makes access problematic. In neuroscience, the power of studying combinatorially defined neuronal populations increases significantly when multiple distinct neuronal features, such as birthdate, collateral projection targets, molecular profiles, or a functional requirement in a given physiological process or behavior can be examined in parallel to reveal deeper insights into the mechanisms involved in the establishment and maintenance of neural circuit organization. However, the ability to carry out such studies is restricted by the limited number of publicly available intersectional mouse lines as well as the inherent difficulty in developing new intersectional alleles.

We present a novel CRISPR/Cas9-mediated system for consistently targeting large cassettes to the *Rosa26* locus for high throughput and parallel generation of intersectional and conditional mouse lines. These studies demonstrate that CRISPR/Cas9 can be used to increase targeting efficiency at the *Rosa26* locus in mouse embryonic stem cells with shortened 1-1.2 kb homology arms while preserving pluripotency and the ability to transmit alleles via the germline. This technology allows for the opportunity to generate mouse lines that express a wide range of effector molecules in various cell types, enabling the developmental, anatomical, molecular, and functional characterization of cellular organization in behavior and physiology. To democratize the generation of intersectional and conditional targeted ES cells and mouse lines, we have generated a publicly available vector toolkit for single-step cloning into *Rosa26-*targeting vectors for double and single-recombinase responsive cassettes (**Table 1).** We have also generated several mouse lines available, without restriction, to the not-for-profit research community enabling high specificity intersectional access of neuronal and other populations for cross correlative mapping approaches using functional neuronal perturbation, molecular profiling, and anatomical characterizations for multifaceted studies (**Table 2**). Overall, these resources and the modular nature of the intersectional approach make rapid and low-cost production of large numbers of intersectional alleles possible that can be efficiently distributed as ES cells or mouse lines throughout the mouse research community. All vectors (Addgene; 97007-97012, 99142), mouse lines (MMRRC; 043513-043519), and ES cells (BCM ES Cell Core Facility) have been made publicly and unconditionally available to the academic research community.

## METHODS

### Construction of targeting and Cas9 vectors

The simplified *Rosa26* targeting vector was derived using standard cloning procedures from a targeting vector used in previous studies (52) consisting of a p15a origin of replication, kanamycin selection cassette, DTA negative selection cassette, and *Rosa26* homology (1081 bp 5’ homology arm and 4342 bp 3’ homology arm). The total vector size was 9084 bps. The DTA was removed and the 5’ homology arm was shortened to 1025 bp fragment (deleting the sgRNA target sequence (chr6:113,076,062-113,077,087 GRCm38/mm10)) while the 3’ homology arm was shortened to 1231 bps (chr6:113,074,801-113,076,032 GRCm38/mm10), maintaining the insertion site within the first intron and reducing the total vector length to 4694 bps and eliminating a number of repetitive regions that often resulted in vector instability when constructing intersectional targeting vectors.

For the Cas9 expressing vector, we selected an sgRNA target sequence that was close to the 5’ and 3’ junction of the *Rosa26* gene locus (ACTGGAGTTGCAGATCACGA with PAM motif GGG; chr6:113,076,040-113,076,05 GRCm38/mm10). The selected sgRNA was cloned into the BbsI sites of the px330 vector that expresses Cas9 (final vector is called *px330_Rosa26_sgRNA*).

Knock-in cassettes were assembled in either 1) a previously used intersectional template plasmid (*RR1-4, RR6*-*7*); 2) a newly constructed intersectional template plasmid with neomycin promoter and pA elements removed (*RR5* and *RR8*); or 3) a constructed Dre-responsive template plasmid (*RR9* and *RR10*). The intersectional plasmid consisted of a ubiquitous *CAG* promoter sequence; an *FRT*-flanked stop cassette consisting of a PGK-neo sequence (for positive selection of targeted ES cells) and three *SV40 pA* sequences; a *LoxP*-flanked stop cassette containing mCherry and a PBS302 stop cassette; and a cloning site for insertion followed by a *WPRE* and *BGHpA* sequence. The new intersectional template consisted of a ubiquitous *CAG* promoter sequence; an *FRT* flanked stop cassette consisting of a neomycin sequence and *His3_SV40 pA*; a *LoxP*-flanked subtractive cloning site stop cassette consisting of a *His3_SV40 pA*; and a cloning site for insertion followed by *WPRE* and *BGHpA* sequences. The Dre-responsive template consisted of a *Rox*-flanked stop cassette consisting of a neomycin sequence and PBS302 stop cassette, and a cloning site followed by a *WPRE* and *BGHpA* sequence.

For assembly of complete targeting vectors, the cDNAs of interest were PCR amplified and cloned into the corresponding vector. The intersectional or Dre-responsive cassettes were then cut out with PacI or PacI/AscI and cloned into the shortened *Rosa26* targeting vector.

To facilitate single step cloning of complete targeting vectors, we generated a vector toolkit (**Table 1)** by constructing and moving empty *CAG* cassettes in the Rosa26 targeting vector. We generated four template vectors with neomycin resistance: 1) a Cre/FLP responsive targeting vector; 2) Cre only; 3) FLP only; and 4) Dre-responsive targeting vector. The intersectional template vector consists of the *CAG* promoter, an *FRT*-flanked neomycin resistance gene and stop cassette, a subtractive cloning site (EcoRV) and stop cassette flanked by *LoxP* sites, an intersectional cloning site (PmeI), and *WPRE* and *BHGpA* elements. Each single template vector consists of the *CAG* promoter, stop cassette(s) and neomycin resistance cassette flanked by recombinase sites, cloning site (SwaI), and *WPRE* and *bgh polyA* elements. Additionally, for possible direct oocyte injections we also generated a Cre/FLP responsive targeting vector that does not contain a neomycin resistance gene, with a subtractive cloning site (EcoRV) and intersectional cloning site (SwaI). All cloning sites are blunt restriction enzyme sites for cloning facilitation.

Sequences of the template vectors and *px330* vector and plasmids are available through Addgene (provisional ID #s 97007-97012, 99142).

### Generation of knock-in mice

Embryonic stem (ES) cells (AB2.2) were electroporated with 15-20 μg of varying ratios of the *px330_Rosa26_sgRNA* vector to the targeting vector. Neomycin selected clones were screened for homologous recombination using PCR genotyping. Targeted clones were identified using PCR genotyping for 5’ and 3’ targeting from outside the homology arm and were considered to be successful targeting events if positive for both 5’ and 3’ genotyping. We used pairs 5’CGCCTAAAGAAGAGGCTGTG (Rosa26-F) and 5’GGGCGTACTTGGCATATGAT (CAG-R), producing a 1450 bp band; and 5’AATCAACCTCTGGATTACAAAATTT (WPRE-F) and 5’TGGCTCCTCTGTCCACAGTT (Rosa26-R), producing a 2472 bp band.

Select targeted clones were microinjected into C57B1/J6 blastocysts and chimeric males were bred to wildtype C57B1/J6 females to achieve germline transmission.

### Pronuclear injection

We constructed a targeting vector by cloning a calcium indicator, Twitch-2B (55), into the Cre responsive *Rosa26* targeting vector. The Baylor College of Medicine Embryonic Stem (ES) Cell Core generated the *Rosa26*-specific sgRNA and Cas9 protein for injection. Pronuclear injections were performed by the Baylor College of Medicine Genetically Engineered Mouse (GEM) Core using the following parameters: 30 ng/μl Cas9 protein, 20 ng/μl sgRNA, and 2 ng/μl dsDNA plasmid. Potential founders were screened for targeting as described above.

### Breeding, genetic background, and maintenance of mice

We maintained colonies of select mouse strains by backcrossing to C57BL/6J mice. For routine genotyping, we carried out PCR amplification of DNA from ear punch preparations using the boiling alkaline lysis procedure. *Rosa26* specific primers for the mice were 5’-GCACTTGCTCTCCCAAAGTC, 5’-GGGCGTACTTGGCATATGAT, and 5’-CTTTAAGCCTGCCCAGAAGA, and yield a 495 bp band (targeted) and 330 bp band (wt). For histology experiments, *RR5* mice were bred to *DBH^p2aFLPo^*(54)*; B6N.FVB-Tmem163^Tg(ACTB-cre)2Mrt^/CjDswJ (Bactin-Cre)* and *DBH^p2aFLPo^* mice. For plethysmography experiments, *B6.FVB(Cg)-Tg(Dbh-Cre)KH212Gsat/Mmucd (DBH-Cre)* mice were mated with homozygous Cre-only responsive *RR1P* and *RR2P* mice (after crossing intersectional alleles to *B6;SJL-Tg(ACTFLPe)9205Dym/J* [JAX 003800] mice to recombine out the FLP-responsive stop cassette followed by homozygosity) to derive animals in which all mice carried the Cre-responsive hM4D or hM3D allele. Sibling animals that did not inherit the *Cre* allele were used as control animals to the *Cre* positive offspring. For electrophysiology experiments, *DBH-Cre; RR2P* mice were mated to *RC::ePe* mice expressing a floxed eGFP. Cre-specific primers were 5’-ATCGCCATCTTCCAGCAGGCGCACCATTGCCC and 5’-GCATTTCTGGGGATTGCTTA and yielded a 550 bp band if positive. FLPo-specific primers were 5’CACGCCCAGGTACTTGTTCT and 5’CCACAGCAAGAAGATGCTGA and yielded a 226 bp band if positive.

Established mouse lines reside at the Mutant Mouse Regional Resource Center (MMRRC) and will be unconditionally available to the academic research community (MMRRC 043513-043519). Targeted ES cells have been archived and are available upon request under the condition that any new mouse lines are deposited in a public repository.

All animal experiments were performed with the approval of IACUC.

### Off-target analysis

Off-target sequences were identified using the Optimized CRISPR Design tool (crispr.mit.edu). The top 5 sequences for each sgRNA were amplified from selected targeted ES cells and sequenced to determine if any mutations or changes occurred in off-target sites.

### Viral injection

To test the functionality of the *RR5* mouse strain, adult *RR5* mice were injected with equal titers of AAV9-hSyn-FLPo and AAV1-hSyn-Cre viruses (obtained from M. Xue at BCM, 250 nl at 1.12×10^12^ GC/mL) into the dentate gyrus (coordinates from bregma AP=-2.70, DV=-2.12, ML=1.84) and with equal titers of AAV-EF1a-Cre-WPRE and AAV-EF1a-FLPo-WPRE (UNC Vector Core, 250 nl at 5.06×10^11^ GC/mL) into the amygdala (coordinates from bregma AP=-1.06, DV=-4.61, ML=-2.86) and allowed to incubate for 7-14 days. For expression in the olfactory bulb, adult *RR5* mice were injected with AAV-CAG-Cre (obtained from Neuroconnectivity Core at Jan and Dan Duncan Neurological Research Institute, AAV2/9 690 μL at 5.37×10^13^ pp/mL) and AAV-Ef1a-Flp (obtained from Neuroconnectivity Core at Jan and Dan Duncan Neurological Research Institute, AAV2/9 690 μL at 7.16×10^11^ pp/mL) into the core of the olfactory bulb (coordinates from bregma AP=4.5mm, DV=-2.25mm, ML=±0.8mm) and allowed to incubate for 14 days. For the CAV2-Cre virus experiments, *RR5; DBH^p2aFLP^* mice were injected with CAV2-Cre virus (IGMM Viral Core, 500 nL at 6.0×10^12^ pp/mL) into the amygdala (coordinates -2.86, -1.06, -4.61) and allowed to incubate for 17 days. For the Retro-AAV-Ef1a-FLPo virus experiments, *RR5; Vglut2_Cre* mice were injected with Retro-FLPo virus (obtained from Neuroconnectivity Core at Jan and Dan Duncan Neurological Research Institute, 690 μL at 1.49×10^12^ pp/mL) into the lateral hypothalamic area (coordinates from bregma AP=-1.22mm, DV=-5.12mm, ML=±0.97mm) and allowed to incubate for 21 days.

### Histology

For *RR5* expression, animals were sacrificed and transcardially perfused with 0.1M phosphate-buffered saline (PBS) then with 4% paraformaldehyde (PFA) in PBS. Brains were dissected out and drop fixed for two hours in 4% PFA before a PBS rinse and equilibration in 20% sucrose in PBS. Brains were sectioned into 30-40 μm coronal sections and mounted on slides. Images were collected on a Zeiss confocal LSM780 microscope or Leica TCS SPE confocal microscope.

### Electrophysiology

#### Slice preparation

Slice preparation from the mouse LC follows an N-Methyl-D-glucamine (NMDG) slicing protocol (89, 90). Briefly, animals were deeply anesthetized using 3% isoflurane. After decapitation, the brain was removed and placed into cold (0−4 °C) oxygenated NMDG solution containing 93 mM NMDG, 93 mM HCl, 2.5 mM KCl, 1.2 mM NaH_2_PO_4_, 30 mM NaHCO_3_, 20 mM HEPES, 25 mM glucose, 5 mM sodium ascorbate, 2 mM Thiourea, 3 mM sodium pyruvate, 10mM MgSO_4_ and 0.5 mM CaCl_2_, pH 7.35 (all from SIGMA-ALDRICH). Horizontal slices were prepared using a vibratome (200 µm thick) using zirconia blades. The brain slices were kept at 37.0 ± 0.5 °C in oxygenated NMDG solution for 10 minutes. They were then transferred to an artificial cerebrospinal fluid (ACSF) containing 125 mM NaCl, 2.5 mM KCl, 1.25 mM NaH_2_PO_4_, 25 mM NaHCO_3_, 1 mM MgCl_2_, 25 mM glucose, and 2 mM CaCl_2_ (pH 7.4) for at least 1 hour prior to the beginning of recordings. During the recording sessions, the slices were submerged in a commercially available chamber (Luig Neumann, Order No. 200-100 500 0150-M) and were stabilized with a fine nylon net attached to a custom-designed platinum ring. This recording chamber was continuously perfused with oxygenated physiological solution throughout the recording session.

#### Recordings

Whole-cell recordings were performed as described previously (90–92). Briefly, patch pipettes (2-7 MΩ) were filled with an internal solution containing 120 mM potassium gluconate, 10 mM HEPES, 4 mM KCl, 4 mM MgATP, 0.3 mM Na_3_GTP, 10 mM sodium phosphocreatine and 0.5% biocytin (pH 7.25). Whole-cell recordings from up to 8 LC neurons were performed using two Quadro EPC 10 amplifiers (HEKA Electronic, Germany). PatchMaster (HEKA) and custom-written Matlab-based programs (Mathworks) were used to operate the recording system and perform online and offline data analysis. In current-clamp recordings, neurons were first current clamped at ∼-40pA to prevent spontaneous firing. Prior to investigating the effect of drugs, we calculated spike thresholds and recorded firing patterns in response to sustained depolarizing currents by injecting increasing current steps (+10pA). Continuous recordings were obtained from LC neurons current clamped at -40 to 0 pA during drug wash-on experiments. We also calculated other intrinsic electrophysiological parameters, such as the input resistance, membrane time constant, spike amplitude, after-hyperpolarization (AHP) etc. (90–92).

### Plethysmography

Plethysmography on conscious, unrestrained mice was carried out as described on 6-12 week old adult animals (52, 88). Mice were subjected to a five-day habituation protocol with each day consisting of several minutes of handling, temperature taken by rectal probe, intraperitoneal saline injection, and 30 minutes in the plethysmography chamber. Mice were then tested within one week of the last day of conditioning.

On the day of testing, mice were taken from their home cage, weighed, and rectal temperature was taken. Animals were then placed into an airtight, temperature controlled (∼32**°**C) plethysmography chamber and allowed to acclimate for at least 20 minutes in room air (21% O_2_/79% N_2_) conditions. After acclimation and measurement under room air, the chamber gas was switched to a hypercapnic mixture of 5% CO_2_/21% O_2_/74% N_2_ for 20 minutes. Chamber gas was then switched back to room air for 20 minutes. The mice were briefly removed for rectal temperature measurement and intra-peritoneal injection of clozapine-N-oxide (CNO, National Institute of Mental Health Chemical Synthesis and Drug Supply Program) dissolved in saline (0.1 mg/mL) for an effective concentration 1 mg/kg. The animal was returned to the chamber for another 20 minutes of room air, 20 minutes of hypercapnia, and 20 minutes of room air. The animal was then removed from the chamber and rectal temperature was taken immediately afterwards and again 30 minutes after the termination of the experiment. The animal was placed in its own cage during these 30 minutes at the ambient room temperature (∼23**°**C).

#### Plethysmography data collection and analysis

Plethysmography pressure changes were measured using a Validyne DP45 differential pressure transducer and CD15 carrier demodulator in comparison to a reference chamber and recorded with LabChartPro in real time. Waveforms were analyzed offline using LabChart Pro and custom written MATLAB code (Supplemental Figure 3) to determine respiratory rate (V_f_), tidal volume (V_T_) (52), minute ventilation (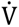_E_), oxygen consumption (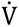_O2_), ventilatory equivalents for oxygen (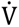_E_/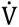_O2_), and pattern analysis. Respiratory waveforms were collected offline during periods when the animal was at rest and readings were free from movement artifacts. A minimum of 1-minute cumulative data compiled from traces at least 10 seconds long from the last 10 minutes of a given experimental condition were analyzed. O_2_ consumption was determined by comparing the gas composition between calibration in an empty chamber and live breathing using an AEI oxygen sensor and analyzer. Chamber temperature was constantly monitored and recorded using a ThermoWorks MicroThermo 2 with probe and was recorded with LabChartPro in real time.

#### Plethysmography statistics

Results (V_f_, V_T_, 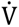_E_, 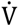_O2_, 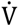_E_/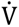_O2_) for room air and hypercapnic data were compared between all cohorts using a linear mixed-effects regression model with animal type (experimental or control) and CNO administration (pre- or post-injection) as fixed effects and animal ID as a random effect. Temperature data was compared using a linear mixed-effects regression model with animal type (experimental or control) as a fixed effect. Residuals were independent and identically distributed as a normal distribution, which matches our model assumptions (Supplemental Figures 4-7). The residual plot for ventilatory equivalents of oxygen in room air for hM4D (Fig. 8) shows a slightly different distribution pattern (Supplemental Figure 5). However, this is because the random effect coefficients are unusually small. This data still follows the normal and independent residuals assumption. Random effect coefficient distribution is not as easily assessed with so few datapoints and is not critical to statistical outcome if violated (93). The p-values reported correspond to statistical significance of the conditional interaction between animal type and CNO administration. A p-value of <0.05 was used to indicate statistical significance and standard error of the mean is shown on all charts.

### Imaging of Twitch 2 in mouse preovulatory follicles

Preovulatory follicles were isolated and imaged as previously described (94). Briefly, antral follicles were dissected from 23- to 26-d-old *RR8; Bact_Cre mice*. Follicles were cultured for 24– 30 h on organotypic membranes (Millipore; cat. No. PICMORG50) in the presence of follicle-stimulating hormone. The follicle was held in a perfusion slide consisting of a plastic slide (ibidi) and a glass coverslip and assembled using silicon grease. The slide was constructed such that medium containing ovine LH (National Hormone and Peptide Program; 10 μg/mL) could be perfused through a 200-μm-deep channel holding the follicle. Temperature was maintained at 30–34 °C, by use of a warm air blower (Nevtek). Preovulatory Follicles were imaged using a Zeiss Pascal confocal microscope with a 40X/1.2 NA objective. Images were collected every 10 seconds. Measurements were corrected for autofluorescence and for spectral bleed-through of CFP into the YFP channel. Ratios were calculated by dividing the mean CFP intensity in each region of interest by the mean YFP intensity. Data analysis was done using ImageJ and Excel software. Data is representative of 4 follicles.

## Declarations

### Ethics approval and consent to participate

All experiments reported herein were conducted with explicit approval of and oversight by both Baylor College of Medicine Institutional Animal Care and Use Committee (IACUC) and University of Connecticut IACUC and abide by all state and national regulations regarding animal research for each work site.

### Consent for publication

Not applicable.

### Availability of data and materials

All datasets, animals, and materials generated and/ or used in this study are publicly available (Addgene and Mutant Mouse Resource and Research Centers Supported by NIH (MMRRC)) or available from the corresponding author on reasonable request.

### Funding

R01: H1130249 (RR)

R21: OD025327 (RR)

R37: HD014939 (JRE and Laurinda A. Jaffe)

### Authors’ contributions

SJL completed experiments, analyzed data, and prepared the manuscript.

AM completed experiments, analyzed data, and prepared the manuscript.

PJH completed experiments, analyzed data, and prepared the manuscript.

PGF completed experiments, analyzed data and prepared the manuscript.

JP completed experiments, analyzed data, and prepared the manuscript.

AC completed statistical analyses and interpretations and prepared the manuscript.

JJS completed experiments, analyzed data, and prepared the manuscript.

VKM assisted in the completion of experiments.

PJZ completed experiments, analyzed data and prepared the manuscript

JRE completed experiments, analyzed data, and prepared the manuscript.

GA statistical analyses and interpretations and prepared the manuscript.

XJ completed experiments, analyzed data, and prepared the manuscript.

BRA completed experiments, analyzed data, and prepared the manuscript.

AST completed experiments, analyzed data and prepared the manuscript

M C-M assisted with experimental design and prepared the manuscript

RR conceptualized the study, performed experiments, analyzed data, and prepared the manuscript.

All authors read and approved the final manuscript.

## Acknowledgements

We thank M. Xue for providing the Cre and FLPo AAV viruses used in the dentate gyrus injection and Ronda Kram for technical assistance with cloning, mouse husbandry, and maintenance. We thank J. Dougherty, E. Schuman, J. Wess, B. Roth, and S. Sternson for providing plasmid templates for the EGFP-L10A, MetRS, G_s_-D, hM4D and hM3D, and PSAM cDNA, respectively. We thank Laurinda A. Jaffe for collaborating with us and author JRE with the Twitch studies. The Baylor College of Medicine Embryonic Stem (ES) Cell Core performed the ES cell electroporations and were exceptionally helpful in our experiments and the Baylor College of Medicine Genetically Engineered Mouse (GEM) Core performed the blastocyst injections. Imaging was carried out at the Baylor College of Medicine Optical Imaging and Vital Microscopy Core.

---

**Supplemental Figure 1. Genetic schema for animals used to gather data for Figures 3-4.**

**Supplemental Figure 2. Genetic schema for animals used in Figures 7-9.**

**Supplemental Figure 3. Custom MatLab code for plethysmography data analysis.**

**Supplemental Table 4. QQplots for normal distribution testing of residuals from plethysmography analyses from Fig. 8.**

**Supplemental Table 5. Residual plots for independence testing of residuals from plethysmography analyses from Fig. 8.**

**Supplemental Table 6. QQplots for normal distribution testing of residuals from plethysmography analyses from Fig. 9.**

**Supplemental Table 7. Residual plots for independence testing of residuals from plethysmography analyses from Fig. 9.**

