## Supplemental Figures and Tables for "A CRISPR toolbox for generating intersectional genetic mice for functional, molecular, and anatomical circuit mapping"

### Supplemental Figure 1

#### A Genetic schema for Figure 3 A-H

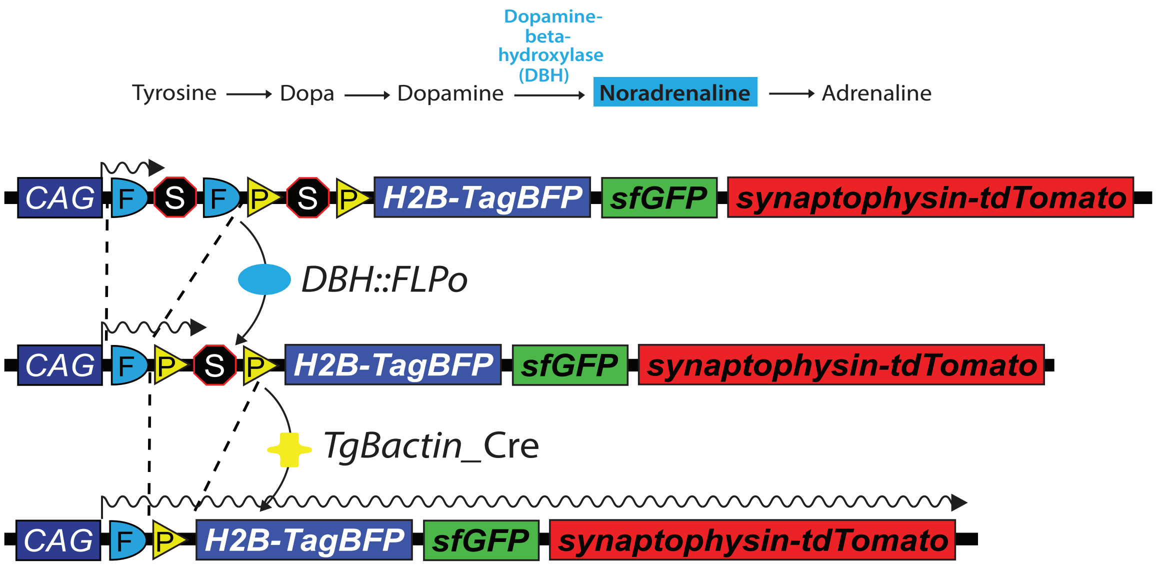


#### B Genetic schema for Figure 3 I-P

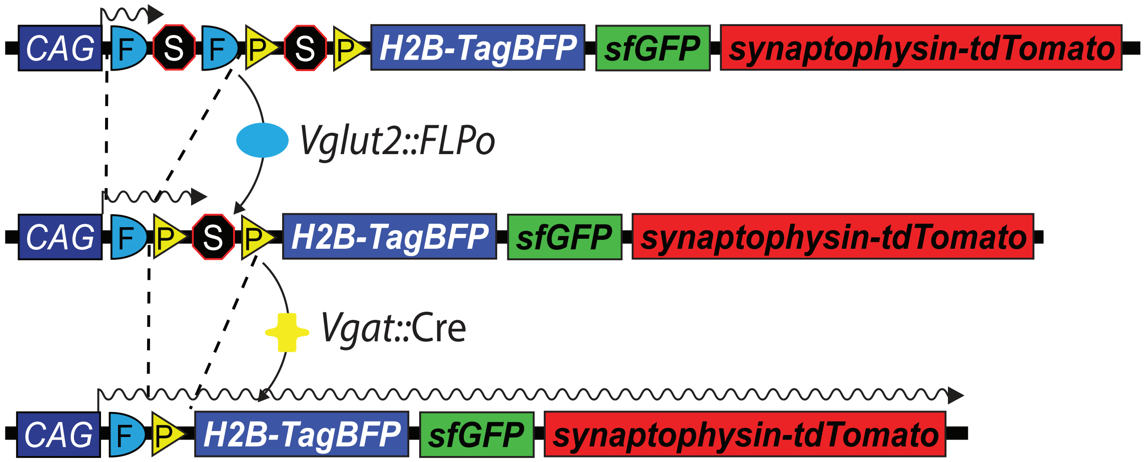


#### C Genetic schema for Figure 4

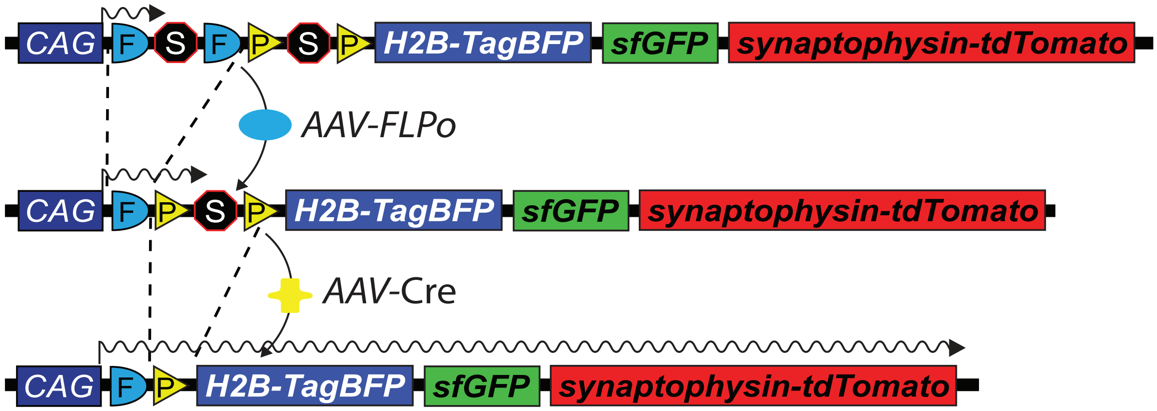

### Supplemental Figure 2

Breeding schema for *RC::P_DREADD; RC::epe; TgDBH_Cre*

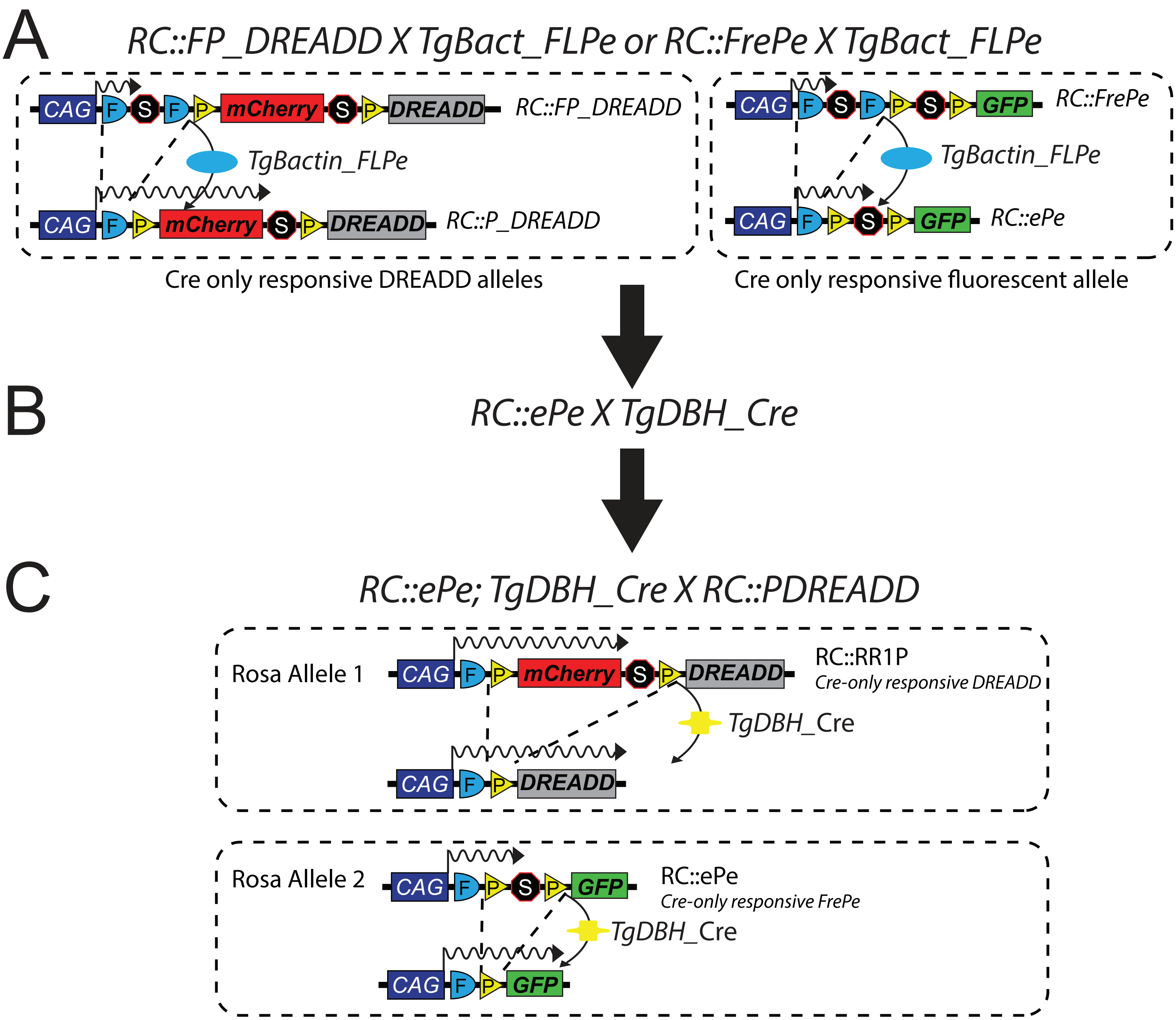

**Supplemental Figure 3** MatLab code for plethysmography analysis.

```
function runStatsHypercapnicCNO(CompiledDataSetRA, CompiledDataSetCO2)

% CompiledDataSetRA is a dataset containing respiratory measurements of each individual animal with
% the following headers for room air conditions:

% Vf | VT | VT(normalized to body weight) | VE | VO2 | VCO2 | VE/VO2 | CNO (Pre or Post) | Condition (RA or CO2) | Animal (Exp or Con) | MUID (Mouse ID #) |


%CompiledDataSetCO2 is a similar dataset containing respiratory measures
% under 5% CO2 conditions

dirname = strcat(pwd, '/Resid');
mkdir(dirname);
% Run models on each of the Room Air Measurements
pValuesRA = cell(3,8);

lmeVf_RA = fitlme(CompiledDataSetRA, 'Vf ~ Animal * CNO + (1|MUID)');
pValuesRA(:,1) = lmeVf_RA.Coefficients.Name(2:4);
pValuesRA(:,2) = num2cell(lmeVf_RA.Coefficients.pValue(2:4));
plotResiduals(lmeVf_RA, 'probability');
saveas(gcf, strcat(dirname, '/lmeVf_RA_qqnorm.png'));
plotResiduals(lmeVf_RA, 'fitted');
saveas(gcf, strcat(dirname, '/lmeVf_RA_resid.png'));

lmeVT_RA = fitlme(CompiledDataSetRA, 'VT ~ Animal * CNO + (1|MUID)');
pValuesRA(:,3) = num2cell(lmeVT_RA.Coefficients.pValue(2:4));
plotResiduals(lmeVT_RA, 'probability');
saveas(gcf, strcat(dirname, '/lmeVT_RA_qqnorm.png'));
plotResiduals(lmeVT_RA, 'fitted');
saveas(gcf, strcat(dirname, '/lmeVT_RA_resid.png'));

lmeVTg_RA = fitlme(CompiledDataSetRA, 'VTg ~ Animal * CNO + (1|MUID)');
pValuesRA(:,4) = num2cell(lmeVTg_RA.Coefficients.pValue(2:4));
plotResiduals(lmeVTg_RA, 'probability');
saveas(gcf, strcat(dirname, '/lmeVTg_RA_qqnorm.png'));
plotResiduals(lmeVTg_RA, 'fitted');
saveas(gcf, strcat(dirname, '/lmeVTg_RA_resid.png'));

lmeVE_RA = fitlme(CompiledDataSetRA, 'VE ~ Animal * CNO + (1|MUID)');
pValuesRA(:,5) = num2cell(lmeVE_RA.Coefficients.pValue(2:4));
plotResiduals(lmeVE_RA, 'probability');
saveas(gcf, strcat(dirname, '/lmeVE_RA_qqnorm.png'));
plotResiduals(lmeVE_RA, 'fitted');
saveas(gcf, strcat(dirname, '/lmeVE_RA_resid.png'));

lmeVO2_RA = fitlme(CompiledDataSetRA, 'VO2 ~ Animal * CNO + (1|MUID)');
pValuesRA(:,6) = num2cell(lmeVO2_RA.Coefficients.pValue(2:4));
plotResiduals(lmeVO2_RA, 'probability');
saveas(gcf, strcat(dirname, '/lmeVO2_RA_qqnorm.png'));
plotResiduals(lmeVO2_RA, 'fitted');
saveas(gcf, strcat(dirname, '/lmeVO2_RA_resid.png'));

lmeVCO2_RA = fitlme(CompiledDataSetRA, 'VCO2 ~ Animal * CNO + (1|MUID)');
pValuesRA(:,7) = num2cell(lmeVCO2_RA.Coefficients.pValue(2:4));
plotResiduals(lmeVCO2_RA, 'probability');
saveas(gcf, strcat(dirname, '/lmeVCO2_RA_qqnorm.png'));
plotResiduals(lmeVCO2_RA, 'fitted');
saveas(gcf, strcat(dirname, '/lmeVCO2_RA_resid.png'));

lmeVEVO2_RA = fitlme(CompiledDataSetRA, 'VEVO2 ~ Animal * CNO + (1|MUID)');
pValuesRA(:,8) = num2cell(lmeVEVO2_RA.Coefficients.pValue(2:4));
plotResiduals(lmeVEVO2_RA, 'probability');
saveas(gcf, strcat(dirname, '/lmeVEVO2_RA_qqnorm.png'));
plotResiduals(lmeVEVO2_RA, 'fitted');
saveas(gcf, strcat(dirname, '/lmeVEVO2_RA_resid.png'));

titlesRA = {'Room Air' 'Vf' 'VT' 'VTg' 'VE' 'VO2' 'VCO2' 'VEVO2'};
pValuesRA = [titlesRA; pValuesRA];

% Present a table with p-values of each test

%f = figure('Position',[100 100 650 120]);
%t = uitable(f,'Data',pValuesRA,'ColumnWidth',{75},'Position',[10 4 1500 102]);

% Run linear mixed-effects regression model on each of the CO2 measurements

pValuesCO2 = cell(3,8);
lmeVf_CO2 = fitlme(CompiledDataSetCO2, 'Vf ~ Animal * CNO + (1|MUID)');
pValuesCO2(:,1) = lmeVf_CO2.Coefficients.Name(2:4);
pValuesCO2(:,2) = num2cell(lmeVf_CO2.Coefficients.pValue(2:4));
plotResiduals(lmeVf_CO2, 'probability');
saveas(gcf, strcat(dirname, '/lmeVf_CO2_qqnorm.png'));
plotResiduals(lmeVf_CO2, 'fitted');
saveas(gcf, strcat(dirname, '/lmeVf_CO2_resid.png'));

lmeVT_CO2 = fitlme(CompiledDataSetCO2, 'VT ~ Animal * CNO + (1|MUID)');
pValuesCO2(:,3) = num2cell(lmeVT_CO2.Coefficients.pValue(2:4));
plotResiduals(lmeVT_CO2, 'probability');
saveas(gcf, strcat(dirname, '/lmeVT_CO2_qqnorm.png'));
plotResiduals(lmeVT_CO2, 'fitted');
saveas(gcf, strcat(dirname, '/lmeVT_CO2_resid.png'));

lmeVTg_CO2 = fitlme(CompiledDataSetCO2, 'VTg ~ Animal * CNO + (1|MUID)');
pValuesCO2(:,4) = num2cell(lmeVTg_CO2.Coefficients.pValue(2:4));
plotResiduals(lmeVTg_CO2, 'probability');
saveas(gcf, strcat(dirname, '/lmeVTg_CO2_qqnorm.png'));
plotResiduals(lmeVTg_CO2, 'fitted');
saveas(gcf, strcat(dirname, '/lmeVTg_CO2_resid.png'));

lmeVE_CO2 = fitlme(CompiledDataSetCO2, 'VE ~ Animal * CNO + (1|MUID)');
pValuesCO2(:,5) = num2cell(lmeVE_CO2.Coefficients.pValue(2:4));
plotResiduals(lmeVE_CO2, 'probability');
saveas(gcf, strcat(dirname, '/lmeVE_CO2_qqnorm.png'));
plotResiduals(lmeVE_CO2, 'fitted');
saveas(gcf, strcat(dirname, '/lmeVE_CO2_resid.png'));

lmeVO2_CO2 = fitlme(CompiledDataSetCO2, 'VO2 ~ Animal * CNO + (1|MUID)');
pValuesCO2(:,6) = num2cell(lmeVO2_CO2.Coefficients.pValue(2:4));
plotResiduals(lmeVO2_CO2, 'probability');
saveas(gcf, strcat(dirname, '/lmeVO2_CO2_qqnorm.png'));
plotResiduals(lmeVO2_CO2, 'fitted');
saveas(gcf, strcat(dirname, '/lmeVO2_CO2_resid.png'));

lmeVCO2_CO2 = fitlme(CompiledDataSetCO2, 'VCO2 ~ Animal * CNO + (1|MUID)');
pValuesCO2(:,7) = num2cell(lmeVCO2_CO2.Coefficients.pValue(2:4));
plotResiduals(lmeVCO2_CO2, 'probability');
saveas(gcf, strcat(dirname, '/lmeVCO2_CO2_qqnorm.png'));
plotResiduals(lmeVCO2_CO2, 'fitted');
saveas(gcf, strcat(dirname, '/lmeVCO2_CO2_resid.png'));

lmeVEVO2_CO2 = fitlme(CompiledDataSetCO2, 'VEVO2 ~ Animal * CNO + (1|MUID)');
pValuesCO2(:,8) = num2cell(lmeVEVO2_CO2.Coefficients.pValue(2:4));
plotResiduals(lmeVEVO2_CO2, 'probability');
saveas(gcf, strcat(dirname, '/lmeVEVO2_CO2_qqnorm.png'));
plotResiduals(lmeVEVO2_CO2, 'fitted');
saveas(gcf, strcat(dirname, '/lmeVEVO2_CO2_resid.png'));

titlesCO2 = {'5% CO2' 'Vf' 'VT' 'VTg' 'VE' 'VO2' 'VCO2' 'VEVO2'};
pValuesCO2 = [titlesCO2; pValuesCO2];

% Present a table with p-values of each test
%f2 = figure('Position',[100 100 650 120]);
%t = uitable(f2,'Data',pValuesCO2,'ColumnWidth',{75},'Position',[10 4 1500 102]);

% Save statistical data
Directory = pwd;
FileName = 'Stats';
FullFileName = fullfile(Directory,FileName);
save(FullFileName);
```

### Supplemental Figure 4

#### QQplots for normal distribution testing of residuals from plethysmography analyses from Fig. 8, RR1 (hM4D).

Room Air | 5% $CO_2$

Respiratory Rate ($V_f$)

Tidal Volume ($V_T$)

Minute Ventilation ($V_E$)

Oxygen Consumption ($V_{O2}$)

Ventilatory Equivalents of Oxygen ($V_E$/$V_{O2}$)

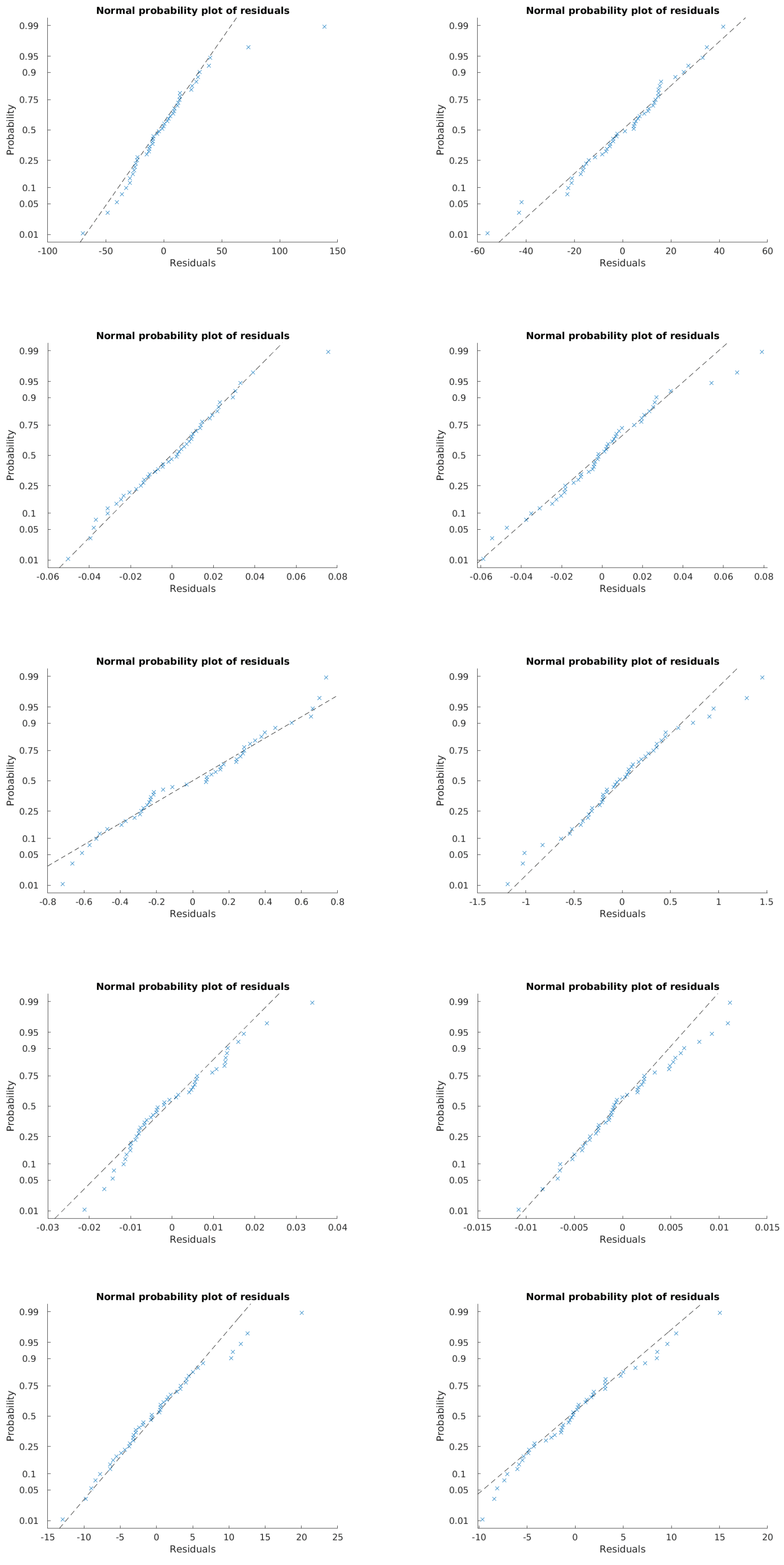

### Supplemental Figure 5

#### Residual plots for independence testing of residuals from plethysmography analyses from Fig. 8, RR1 (hM4D).

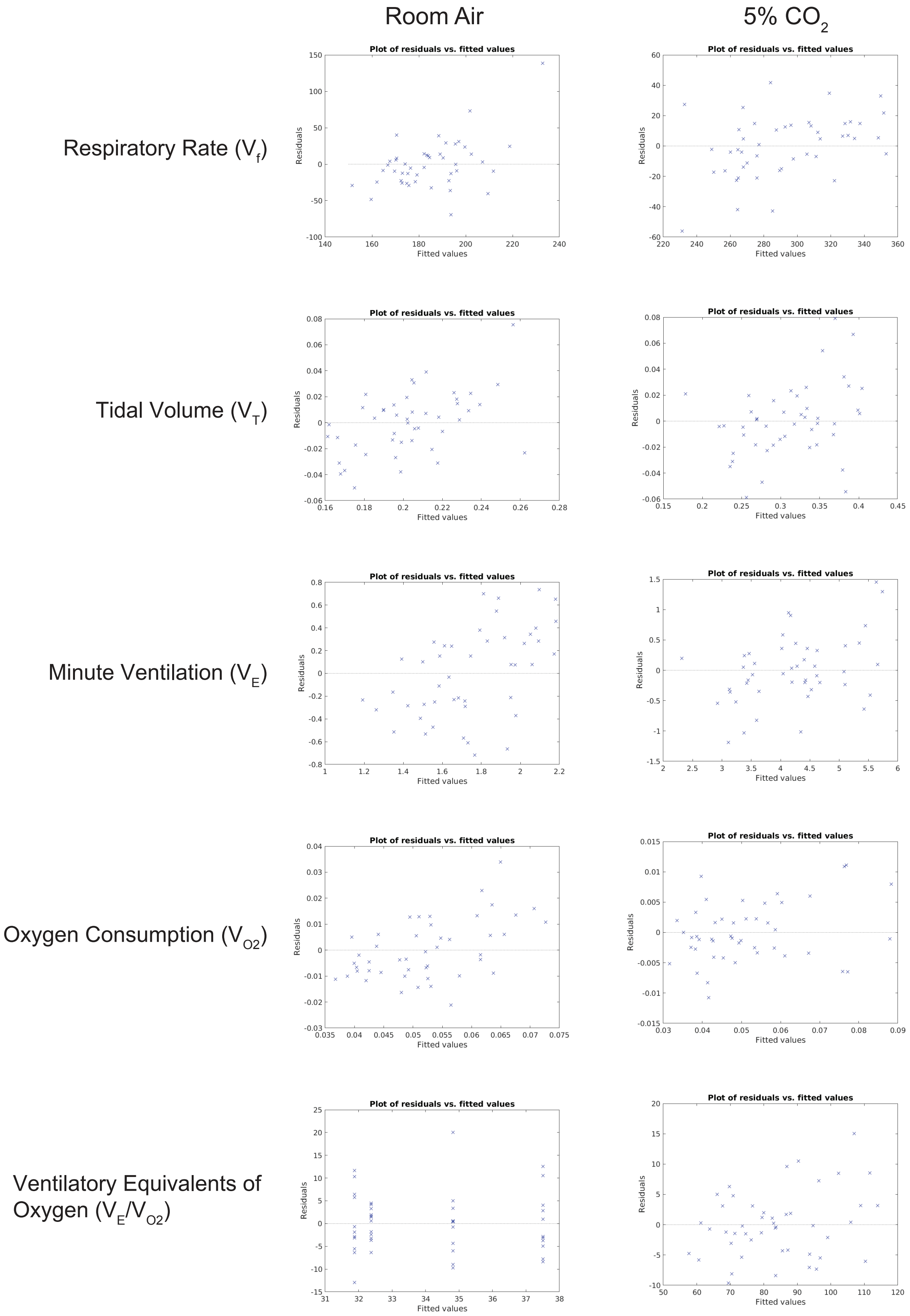

### Supplemental Figure 6

#### QQplots for normal distribution testing of residuals from plethysmography analyses from Fig. 9, RR2 (hM3D).

| | Room Air | 5% $CO_2$ |
|---|---|---|

Respiratory Rate ($V_f$)

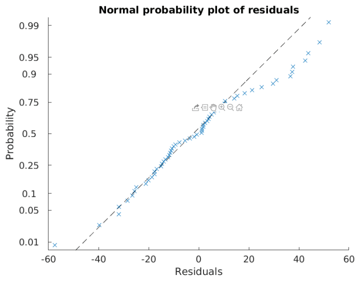


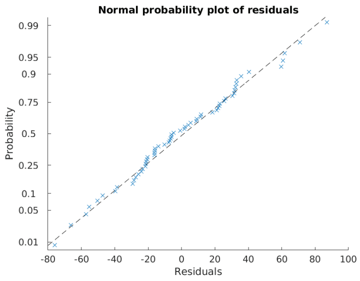


Tidal Volume ($V_T$)

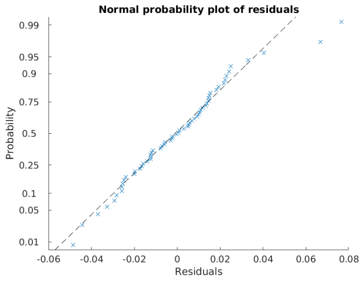


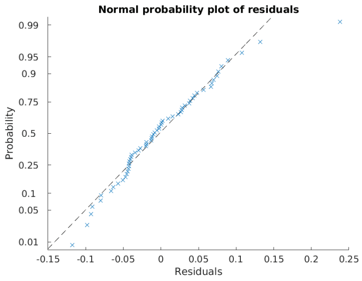


Minute Ventilation ($V_E$)

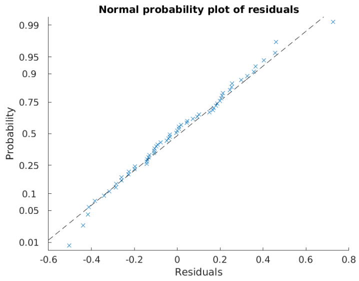


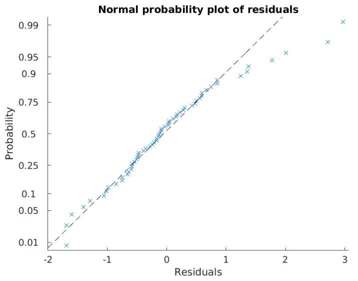


Oxygen Consumption ($V_{O2}$)

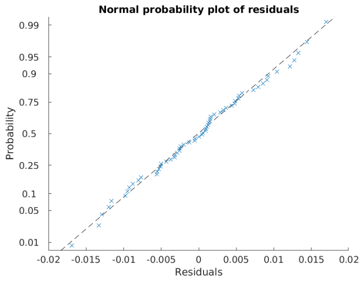


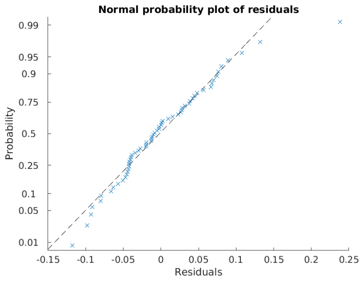


Ventilatory Equivalents of Oxygen ($V_E/V_{O2}$)

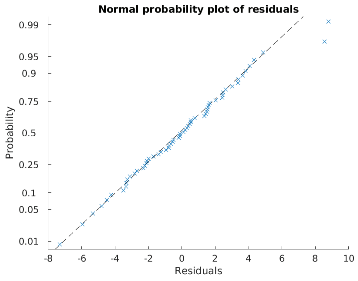


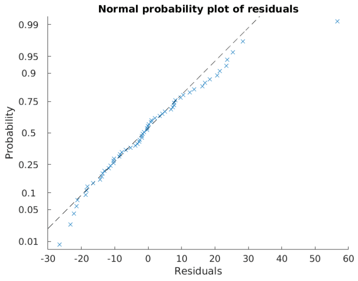

### Supplemental Figure 7

Residual plots for independence testing of residuals from plethysmography analyses from Fig. 9, RR2 (hM3D).

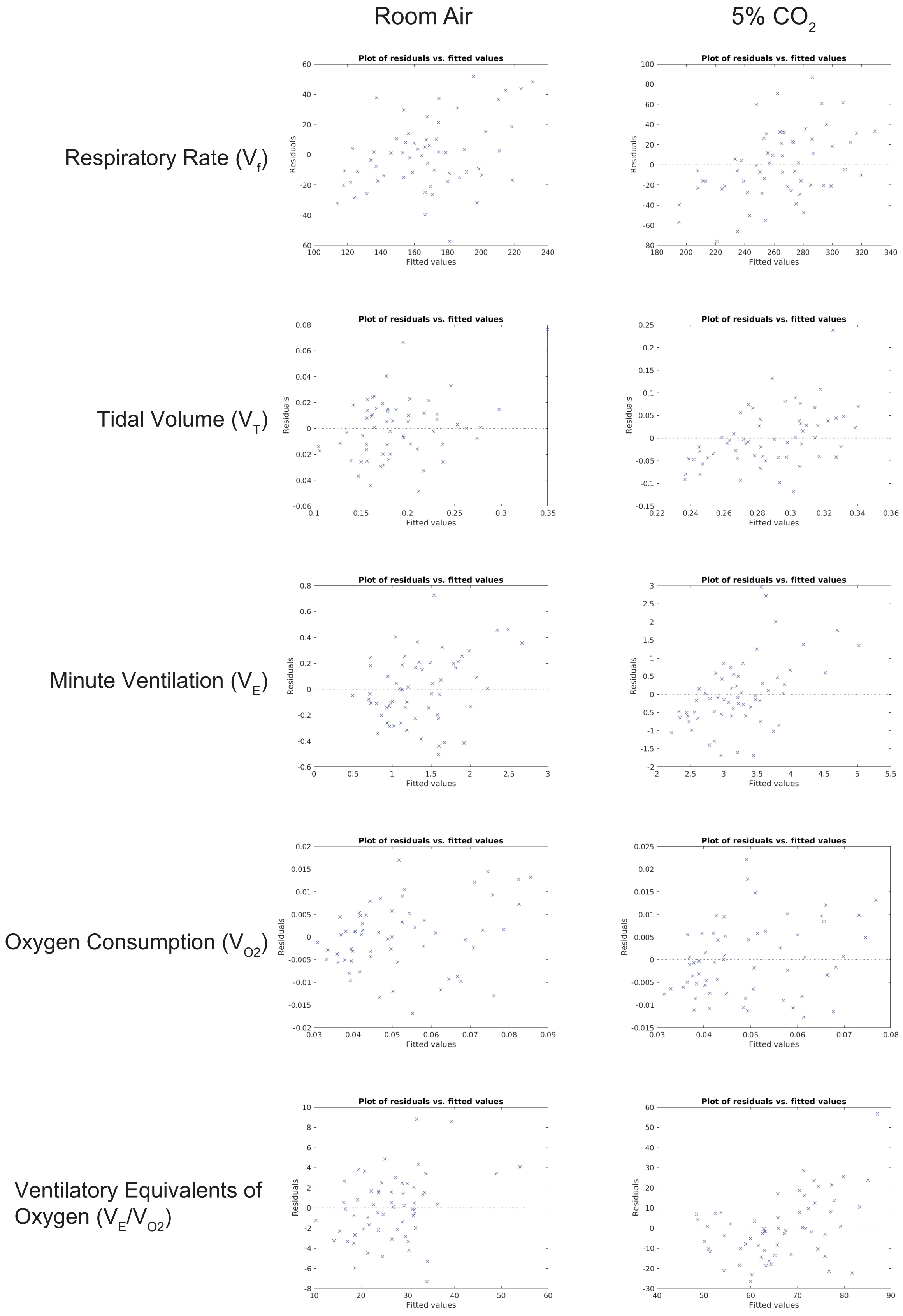
